# Bouts of rest and physical activity in C57BL/6J mice

**DOI:** 10.1101/2023.01.05.522835

**Authors:** K. Pernold, E. Rullman, B. Ulfhake

**Affiliations:** Div. clinical physiology, Department of Laboratory medicine, Karolinska Institutet, CA

## Abstract

The objective was to exploit the raw data output from a scalable home cage (type IIL IVC) monitoring (HCM) system (DVC®), to characterize pattern of undisrupted rest and physical activity (PA) of C57BL/6J mice. The system’s tracking algorithm show that mice in isolation spend 67% of the time in bouts of long rest (≥40s) and 59 % of the time was interpreted as sleep. Twenty percent is physical activity (PA), split equally between local movements and locomotion. Decomposition revealed that a day contains ∼6500 discrete bouts of short and long rest, local and locomotor movements. Mice travel ∼330m per day, mainly during the dark hours, while travelling speed is similar through the light-dark cycle. Locomotor bouts are usually <0.2m and <1% are >1m. Tracking revealed also fits of abnormal behaviour. The starting positions of the bouts showed no preference for the rear over the front of the cage floor, while there was a strong bias for the peripheral (75%) over the central floor area. The composition of bouts has a characteristic circadian pattern, however, intrusive husbandry routines increased bout fragmentation by ∼40%.

Extracting electrode activations density (EAD) from the raw data yielded results close to those obtained with the tracking algorithm, with 59% of time in long rest (<1 EAD s^-1^) and 20% in PA. We confirm that EAD correlates closely with movement distance (r_s_ >0.95) and the data agreed in ∼96% of the file time. Thus, albeit EAD being less informative it may serve as a proxy for PA and rest, enabling monitoring group housed mice. The data show that a change in housing density from one to two, and up to three mice had the same effect size on EAD (∼2) with no difference between sexes. The EAD deviated significantly from this stepwise increase with 4 mice per cage, suggesting a crowdedness stress inducing sex specific adaptations.

We conclude that informative metrics on rest and PA can be automatically extracted from the raw data flow in near-real time (< 1 hrs). These metrics relay useful longitudinal information to those that use or care for the animals.

## Introduction

Metrics of rest and activity in every-day life are important measures of well-being and health in humans and animals alike. These metrics are useful to monitor changes through life, impact of lifestyle, disease signature and progression, and responses to environmental conditions. Traditionally, behaviours of laboratory mice have been assessed by snapshots of home cage behaviours or out-of-cage (and everyday life context) testing, these approaches may not substitute well for cumulative unsupervised monitoring of behavioural alteration over time. Efforts to solve this shortcomings dates more than a century back[1], however, due to technical obstacles it was not until the late 20^th^ century that such systems evolved (e.g. [2, 3]) and later became commercially available (e.g. Intellicage by TSE, Metris by Labora, ActiMot by TSE, and Pheno typer by Noldus; see e.g. [4-8]). Still, these systems are lab-bench type of equipment and not possible to integrate in standard holding systems of the [laboratory animals’] vivarium. More recently novel and scalable HCM systems using different monitoring techniques [9, 10] have been developed for automated non-intrusive 24/7 cumulative monitoring of home-cage activity (home-cage monitoring, HCM) e.g. [11, 12] suitable for a vivarium of small rodents. As recently reviewed [13-15], such HCM systems provide an excellent opportunity to collect cumulative unsupervised records of in-cage rest and PA on a large scale. The purpose of this study was to characterize spontaneous in-cage activity and rest across the circadian cycle and cycles of recurrent husbandry routines over multiple weeks to provide base line data on duration and frequency of bouts of rest and [physical] activity (PA), and how the animals use the cage floor as well as rhythmicity of rest and PA. For this purpose, we recorded cumulative data with a home-cage monitoring system (DVC® Tecniplast SpA) of C57BL/6J mice in standard IVC cages (GM500), kept either in isolation (x1) or in groups at different densities (x2, x3, x4). The DVC® system is based on twelve planar capacitance sensing electrodes situated outside and beneath the cage in a standard cage rack. The electrode array defines the spatial resolution and electrode samples are collected at 4 Hz[12, 15, 16]. The 24/7 flow of electrode reads (raw data) is processed to provide spatial and temporal information on electrode activations which is used to delineate bouts of in-cage PA and rest. With animals kept in isolation the data can be used to track the animals’ position and to monitor the animals’ in-cage movements. These capacities of the system have previously been validated towards CCD recordings[16].

In this report we present data on frequency and duration of bouts of rest and PA, across the cage floor, circadian cycle and across weeks of observation. PA bouts are split into bouts of movements on the spot (MOTS)[17, 18] and bouts of locomotion based on distance made during the bout[17, 19]. Bouts of rest were divided into short and long (≥40s) because long bouts of rest correlates closely with sleep[20-24]. Furthermore, we used electrode activation density (EAD)[12, 16] do assess the impact on in-cage synchronized-rest and PA of male and female mice housed at different densities (x1, x2, x3, and x4).

With standard desk top computers, the metrics presented herein can be extracted 24/7 in near-real time (≤ 1h) for all cages in a 60-slot IVC rack using a desktop computer. Since mice mainly are used as models for human conditions in experimental work, this set of information about in-cage life may not only be of value to the those that care or use the mice but may prove to translate well to corresponding assessments made in humans.

## Materials and methods

### Mouse strain, sex, and age

Cohorts of specific pathogen free (SPF, according to FELASAs exclusion list[25, 26]) male (m) and female (f) C57BL/6J mice were delivered by car from Charles River, Germany, (Table 1). Upon arrival subjected to a brief health check, at 6-8 weeks of age mice were randomly allotted to cages and either grouped 2 (x2), 3 (x3), 4 (x4) per cage or kept in isolation (x1).

**Table 1.**
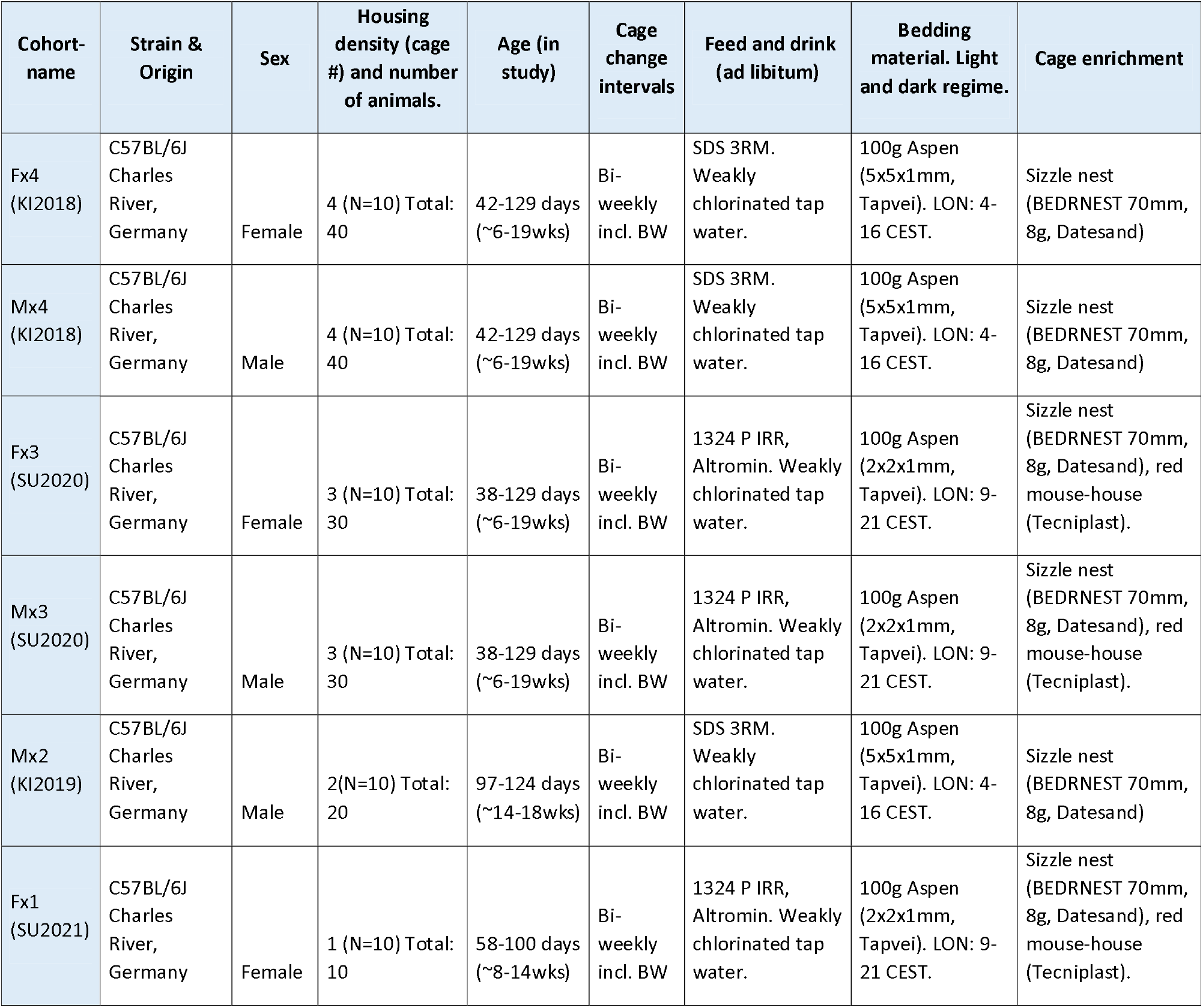

### Holding and husbandry conditions

Mice were kept in individually ventilated caged (IVC) of type GM500 (Tecniplast SpA, Italy) in a DVC system (Tecniplast SpA) (Table 1). IVC cages are ventilated with 75 HEPA14 filtered air exchanges per hour, the air is taken from the holding room and let out through a separate outlet. The holding room has a 12-12h dark/light cycle (DL; Zeitgeber time (ZT) 0-12 (L) and 12-24(D)) with white light level at 15-40 Lux inside the cage. Cohort Fx1, Fx3 and Mx3 had had lights on/ lights off with dawn and dusk of 60 min, while cohorts Mx2, Mx4 and Fx4 had sudden change of the lightning conditions in the vivarium.

All cages had 100g aspen chips 2 or 5 mm (Tapvei, Finland) as bedding, nestlets, Bed-r’Nest or sizzle nest and several (Fx3, Mx3, and Fx1; see Table 1) of the cohorts also had a red polycarbonate mouse house (Tecniplast SpA) as enrichment. The husbandry routines included bi-weekly (see Table 1) cage change (whole cage was changed but red house and some of the soiled beddings were moved along with the animals to the new cage) and also body weighing (Table 1).

Handling of the mice by staff was either by using cupped hand or by forceps at the tail root, all mice in the different groups were subjected to both handling routines. The holding units were subject to health inventories according to FELASA’s recommendation for a sentinel reporting system (i.e. the subjects of the study were not directly affected by the health inventory) four times a year[25, 26] and during the study period the output from the sentinel system used met the FELASA exclusion list for specific pathogen free animals (SPF). Surveillance of health and welfare included daily check-ups and bi-weekly individual examination during the cage-change and weighing. Health is assessed according to a scoring list deployed at all facilities on Karolinska Institutet and Stockholm University, amended by special requirements stated in the ethic permit. When we weighed the animals, these metrics along with the scoring list formed the basis of the welfare and health check-ups. As needed the designated veterinarian of the facility was consulted.

### Ethical considerations

Both husbandry routines and applied procedures followed applicable guidelines and were agreed upon, reviewed, and approved by the Regional Ethics Council, Stockholms Regionala Djurförsöksetiska nämnd; project licenses N116-15, N184/15 plus amendments and project license 9467-2020 with addendum 12337-2021. No special requirements for health and welfare checks beyond those already implemented at the facility (see above) were required by the permits. DVC records of animals kept in isolation derived from 10 cages serving as control animals for an unrelated experiment granted in permit 9467-2020 with addendum 12337-2021.

### DVC recordings

In total, recordings were collected from 60 cages arranged by sex and housing density in 6 cohorts (Table 1) maintained at the Wallenberg Laboratory on Karolinska Institutet or ECF at Stockholm University both in Stockholm, Sweden. Here we have collected new but have also re-analysed previous [27] cumulative DVC recordings for the purpose of revealing patterns of rest and activity. In doing so we reduced number of animals needed for the study.

The core of the DVC system is an electronic sensor board installed externally and below each standard IVC cage of a rack. The sensor board is composed of an array of 12 electrodes and employs a capacitive-based sensing technology (CST). A proximity sensor measures the electrical capacitance of each of the 12 electrodes at 4 Hz (i.e., every 250 msec). The electrical capacitance is influenced by the dielectric properties of matter close to the electrode, leading to measurable capacitance changes due to the presence/ movement of animals in the cage above. Thus, movements across the electrode array are detected and recorded as alterations in capacitance[12, 16]. In this study we used two different analytical approaches based on the CST to reveal pattern of rest and activity in single housed animals.

### REM unit

At the ECF facility, the DVC rack was equipped with a REM unit (Tecniplast SpA) which record 24/7 noise (audible range), vibrations (acceleration), light level (Lux) and presence of humans in front of the rack. Temperature and humidity were regulated by the Scanclime airflow unit modulating these parameters of the air in-flow. At the Wallenberg facility, the air-flow unit uses the air of the holding room, and records of temperature and humidity are those of the holding room. Light level in front of the cages across the DL cycle applicable to cohorts x1 and x3 is shown in Supporting information (Fig. S1).

### Tracking rest and movements

For detailed description of metrics we refer to the manufacturer (tracking) and previous publications (EAD; [12, 16]. Both of these metrics have been validated towards video tracking[16]. Briefly, and as described by the manufacturer the mouse position on the cage floor is determined by estimating a short-period baseline *R*_k_(t) per each electrode *k* and per each time *t* as the maximum capacitance measurement within a 1-minute moving window. *d*_k_(*t*) is the difference between the estimated baseline *R*_k_(*t*) and the current capacitance measurement *c*_k_(*t*) of each electrode. The mouse position is determined as the centroid of the coordinates of the 12 electrodes weighed by their corresponding signal drop *d*_k_(*t*) with a resolution of ∼1mm [15, 16]. A Gaussian filter is applied across time and space to smooth the trajectory.

The tracking algorithm was used to differentiates between the mouse being still (resting), i.e. no change in *x, y* position of the centroid between successive samples, and in motion where *x, y* position change between successive samples (Fig. 1). For practical reasons, cut off point for motion was set when the difference in successive samples of *x, y* [in Euclidian distance] ≥ 1 mm between sample. Based on previous records of step length for this strain and sex, motion-episodes were divided into local movement (movement-on-the-spot, MOTS)[17] being less than one average stride length (65 mm) in radial distance from the starting point and locomotion when the trajectory covered at least one full stride length[19]. Tracking of individual animals is possible only when animals are kept in isolation and were thus executed on cages in cohort Fx1 only (Table 1).

**Fig 1.** An electrode capacitance is calculated from the actual reading of the electrode at t and the base-line value. The values from the 12 electrodes are then used to calculate the x, y position of the mouse centroid (left panel). A PA bout is initiated when x, y changes in successive samples and continuous until the mouse is still again (no change in x, y between successive samples). The time-series of x, y coordinates during a movement bout is used to plot the bout trajectory (right panel) and to decide if it is a MOTS or locomotor bout. Inserted into the track diagram (right panel) is how the cage floor was divided into a front(FF)-rear(RF) area (green line and text) and a central(CF)-peripheral(PF) area of equal size (red line and text), respectively. For further information see text.

The locomotor records covering 6 weeks for each cage, are time-series with bouts of motion (locomotion and MOTS) interrupted by bouts where the animal was still. The duration and frequency of bouts (still, MOTS and locomotion) were calculated, including distance (m) and speed (ms^-1^) along with other metrics when the animal moved. Bouts of being still were subdivided into short and long bouts of rest, based on bout duration. Earlier studies of mice and rats have shown that periods of rest lasting 40s or longer correlate closely with sleep [20-23, 28]. This was used as cut point between long and short rest. The composition of bouts for each cage was compared as aggregated values, light vs dark (night time) hrs., circadian cycle and days, and across weeks, including impact by husbandry routines[12, 16, 27]

### Rest and physical activity (PA) derived from EAD

An alternative to tracking is to extract the spatial and temporal pattern of CST activations as previously described in [12, 16]. With group housed animals rest and PA assessed by EAD does not apply to individual animals only to the group. Rest assessed by EAD is there for referred to as synchronized-rest and, moreover, PA cannot be divided into MOTS and locomotion. In cooperation with the DVC team at Tecniplast, CST activations was extracted from the raw data by taking the average of two consecutive capacitance readings and calculate the difference with the average of the following two readings (two windows, W2) and compared the absolute value of the capacitance change to the lowest possible threshold (λ) that did not pass-through noise generated in empty cages as activity (λ =1.25). The output is binary from the comparison of successive samples, either an electrode is activated (1) or not (0). The method has previously been validated against CCD-tracking[16].

The output was averaged s^-1^ and referred to as electrode activation density s^-1^ (EAD). The average read s^-1^ (average global activation) from all 12 electrodes have been used in most of the analyses of this study. In addition, the density of activations in the front (electrodes 7 to 12) and rear (electrodes 1-6) of the cage was compared to assess polarity of rest and activity, and as a proxy for the spatial extent of PA we computed the number of unique electrodes activated s (UnEA).

Having both the tracking (still-movement) and the EAD (still-PA) records from the animals kept in isolation, allowed us to assess to what extent these two metrics co-variates. The tracking and EAD files for each animal were aligned using the time stamps and compared. Our data confirms previous observations of a close correlation between EAD and tracking distance per hour[16] (Supportive information (Fig. S2)) and the correlation is very close (∼r_s_>0.95; *idem). The corr*elations between distance made during a locomotor or a MOTS bout, on the one hand, and EAD per bout, on the other, revealed that the relationship between made distance and EAD was different between MOTS and locomotor bouts (Supportive information (Fig. S4)) reflecting the different contents of these two bout types. The correlation between locomotor bout distance and EAD was still significant in each animal, however, less close than the cumulative distance per unit time vs. sum EAD per unit time (Supportive information (Fig. S4)). Although the covariation between the EAD and tracking metrics appears to be solid also over an extended period of time (c.f. [16]) and both metrics indicates that animals kept in isolation are at rest on average 77%-80% of the time (see below), there remain some discrepancies when the two data files are compared (Table 2). During 4% of the file time, the tracking coordinates do not change but EADs are recorded (∼18% of all EADs in file across 6 weeks; Table 2). Thus, there is a close but not perfect match between the two metrics.

**Table 2.**
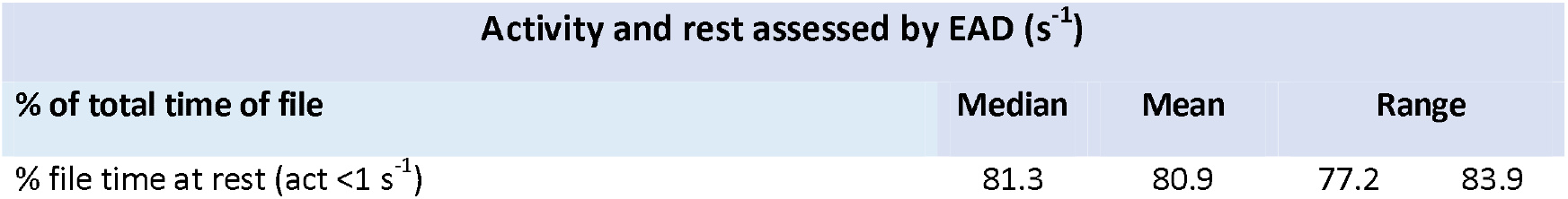
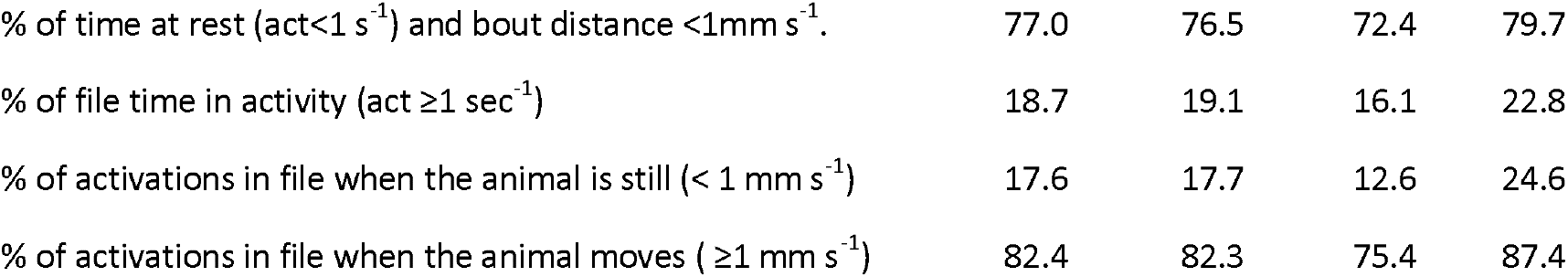

### Data processing

Data were processed through scripts in R (version 4.0.3). The following libraries were used and are hereby acknowledged: broom (ver 0.7.4), compositions (ver 2.0-0), CRamisc (ver 0.5.0.9001), data.table (ver 1.13.6), DescTools (ver 0.99.39), dplyr (ver 1.0.2), filesstrings (ver 3.2.2), ggplot2 (ver 3.3.3), gvlma (ver 1.0.0.3), labelled (ver 2.7.0), microbenchmark (ver 1.4-7), nparLD (ver 2.1), purr (ver 0.3.4), RColorBrewer (ver 1.1-2), readr (ver 1.4.0), stringr (ver 1.4.0), tibble (ver 3.0.4), tidyr (ver 1.1.2), tidyverse (ver 1.3.0), trajar package (ver 1.4.0), zoo (ver 1.8-8), Mixed Effects Models (nlme) library version 3.2.152 on R version 3.5.3.

Time-series of motion and rest (Fx1 cohort only, n=10) and time series of rest and PA of each cage (all cages, n=60; Table 1) generated as described above were used to analyse the distribution of activity and rest by sex and housing density. In animals kept isolated, movement and rest were analysed as aggregated values or segmented into bouts as described above. The data set was then compared to the data set of EAD, i.e. activations and rest [of the same cage], matching the time-series by the time-stamps (s^-1^) to reveal the extent of covariation of metrics among isolated animals (see also [16]). While the locomotion files were divided into bouts of rest and locomotion, and further subdivided into MOTS and locomotion (see above) as well as short and long rest, respectively; the EAD time series files were divided into rest when no electrode was activated (<0.02 average activations s^-1^; i.e., <1 electrode activation s^-1^) and ≥1 electrode activation(s) (≥0.02 activations s^-1^). As with the tracking time-series, episodes (bouts) of activations are intervened by episodes (bouts) of rest. Rest episodes were further subdivided into bouts of short and long rest (≥40s ; see above).

Frequency and duration of bouts were saved along with average and cumulative PA during a bout. The records were used to analyse bout duration and frequency in relation to established rhythmicities [of in-cage behaviours] e.g. day vs. night, circadian, and recurring husbandry routines[10, 12, 29-34].

### Statistical analyses

Aggregated data usually had a normal distribution (Kolmogorov-Smirnoff test of normality) and were analysed by linear-regression, or ANOVA or mixed model ANOVA including post hoc testing. Paired and unpaired samples with a normal distribution were tested with two-sided t-test (equal or unequal variance). Effect size for variables having a normal distribution have been indicated by Hedge’s *q*, or as the coefficient of variation (r^2^)[35]. For parametric statistics we used either R scripts or the plug-in XLSTAT module running on MS Excel. However, several metrics showed large deviations from a normal distribution and could not be normalized by Box-Cox transformation. We therefore choose to test differences across cages, housing densities and/or sexes, cage-change cycles, and days, and across weeks, by nonparametric repeated measures analysis, using the rank-based analysis of variance-type statistic (ATS), as implemented in the nparLD R Software package [36, 37]. Cages are subjects; housing density, time, event, and observation-order are within-subject factors (“sub-plot” repeated factors), and sex and housing density are between-subject factors (“whole-plot” factor) in the models used. The statistical analysis of time-series with nparLD is based on rank-order of the observed data, with the relative effect size (*p*_s_) as effect size measure [37]. The difference towards parametric tests being that instead of the mean difference between observations, the rank-order is used to assess the probability that two sets of observations differ (*p*_s_=0.5 means that there is no difference in rank-order), and/or if the relative effect size varies across the time-series within the sets of observations. Comparison of two independent samples, and paired samples, were conducted using nonparametric Mann-Withney U statistics (U-test and Wilcoxon’s test for matched pairs). In these instances, we used the common language (CL) effect size statistics [38-41]. The CL effect size is based on the rank-order (rank sum) of the observed values and indicates the relative frequency with which the rank sum from one set of observations will be larger than the rank sum of a second set of observations. Correlation between metrics with unknown or a non-normal distribution was done using the nonparametric Spearman rank correlation. The Spearman correlation coefficient (rho, r_s_) indicates the effect size with a range from a perfect inverse covariation (r_s_ =-1), through no covariation (r_s_ =0) to a perfect positive covariation (r_s_ =1) of the ranks for two parameters.

We used R scripts or XLSTAT to run the nonparametric statistical tests. Box plots indicate median, 25%-75% quartiles with max and min as bars. In addition, circular symbols indicate mean values.

## Results

### Tracking of home-cage rest and movements in isolated female mice (Fx1)

For single housed female mice (Fx1, n=10) the total file time of 6 weeks was decomposed into bouts of rest and PA. PA bouts were further segmented in locomotor and MOTS bouts (see Material and methods and [17, 18]). During the observation period, female mice spent 80% of the time at rest with rather small difference between animals (Fig. 2). Sixty-seven percent of the file time were bouts of rest lasting at least 40s. The remaining ∼20% of the time, the animals were engaged in PA split equally between MOTS and bouts of locomotion (*idem)*.

**Fig 2.** Boxplot showing time spent in rest, long rest, short rest, and PA in MOTS and locomotion as fraction of total time. Values indicated are mean, 1st and 3rd quartiles and range. Mean value has been indicated with a blue circle.

### Density and duration of bouts (Fx1 cohort)

A day in life of these ten mice is composed of a string of 4000-8000 bouts (∼0.05-0.1 Hz) of which 18% are rest bouts with median duration of 9s (Table3, Fig 3A-B). Two-point-five percent are long rest bouts with a median duration of 79s (mean duration: 350s; Table 3). Eighty-two percent are bouts of PA (Table 3) and ¾ of these PA bouts are short MOTS bouts (median duration 1.6 s) while 25% are locomotor bouts with a median duration of 5s (Table 3, Fig. 3). Of the daily number of PA bouts, ∼2/3^rds^ occur during lights off when PA bouts are 2.5 more frequent than rest bouts. Thus, of all bouts in file ∼82% are PA bouts with a duration of ≤10s making up <20% of the file time, with MOTS bouts being more than twice as frequent as locomotor bouts (Fig. 3A-B).

**Table 3.**
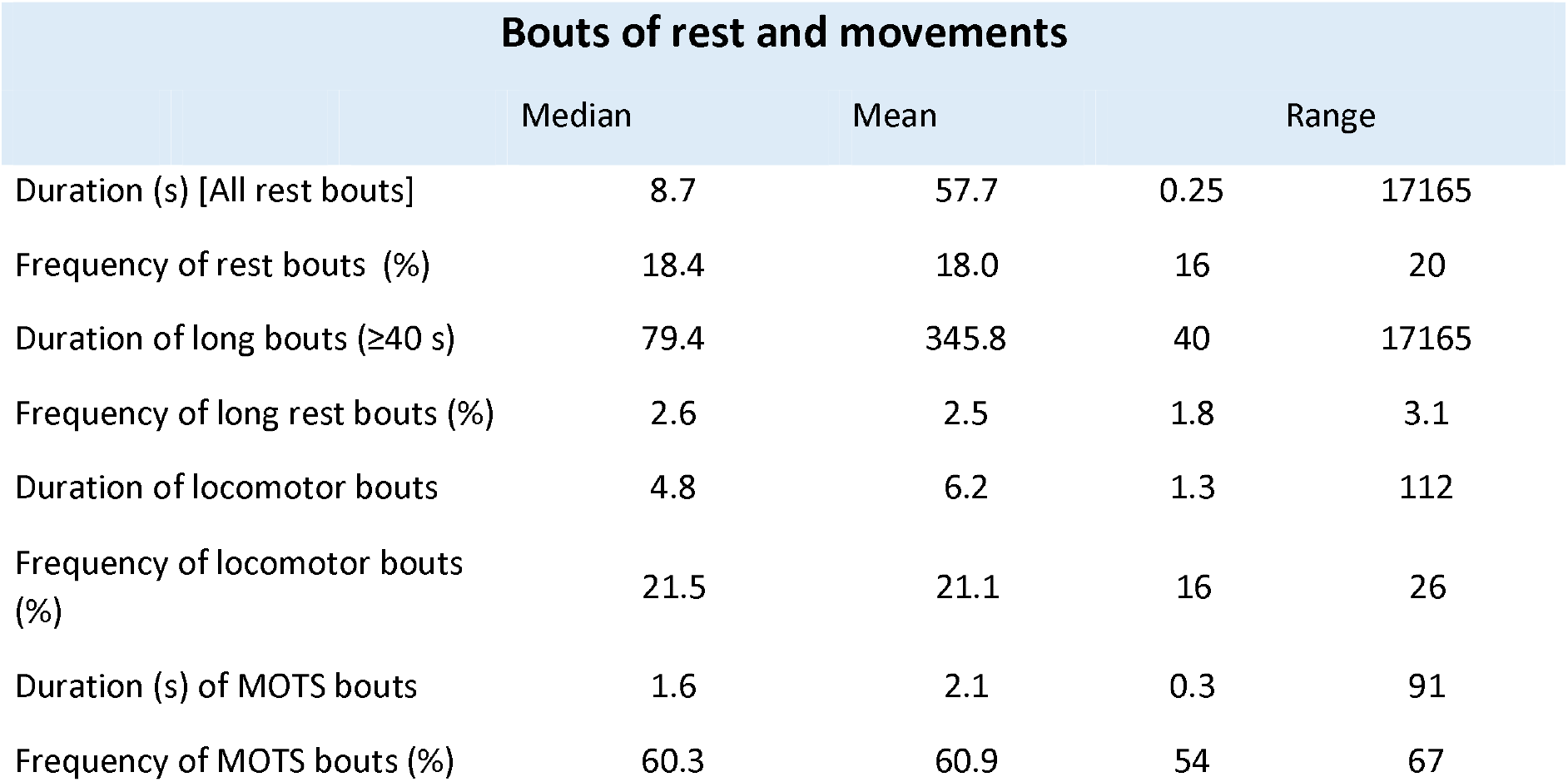

**Fig 3.** (A) Cumulative plot of fraction of the total number of bouts (ordinate) vs bout duration (s, abscissa, logarithmic scale) for all bout types (black line with SD as grey shaded area), rest bouts (black line with blue shaded area indicating SD), MOTS (black line over green area indicating SD), and locomotion bouts (black line on red shaded area indicating SD). (B) Cumulative percentage of total time (ordinate) plotted vs. bout duration (s, abscissa, logarithmic scale). Coloured area and code for bout type are the same as in A. Dotted line in A and B indicates number of bouts (∼90% in A) having a duration <10 s and their combined fraction of total time (∼20% in B).

We confirm previous observations[12, 27] that the time in PA bouts decreases significantly - 30 to 40% (p=2E-9; Fig. 4A) across the cage change cycle, while cycle-to-cycle variation was small (*idem). Converse*ly, the time in rest bouts drops initially followed by an increase by ∼30% (p=2.8E-11; Fig. 4B). The cage change also upsets the pattern of long rest bouts during daytime (lights on) (Fig. 5; see also Supportive information (Fig S6)). Days before a cage-change, long rest usually occurs as 4-6 bouts during day light [intercalated by bouts of PA and short rest] a pattern that is substantially fragmented by this intervention (*idem)*.

**Fig 4.** A and B show impact by day post cage change (dp; abscissa) and cage change cycle (CC 1-3; colour coded grey, red, and green) of fraction of total time spent in bouts of PA (A) and (B) long rest (ordinates). The relative effect size of dp and CC is shown in Supplementary information (Fig. S5). In A (model: PA ∼dp * CC), the major impact is by dp (p=2E-9) with only a minor contribution by CC (p=0.02). B shows that CC has no significant impact on time in long rest (p=0.51) while dp has a strong effect (p=3.8E-11) (model: long rest∼dp*CC).

**Fig 5.** Pattern of long-rest episodes during lights on (ZT 0-12) and lights off (ZT 12-24) (DL 12:12) for the mice S1, S7 and S10 housed in isolation. Long rest bouts indicated by blue colour. Bouts disrupting long-rest periods in black/green. Ordinate is 12h lights on to the left and 12h lights off to the right. Columns are day post cage change (dp) where 0 is the day of the cage change (CC; red arrow indicates the time for CC) for lights on (to the left) and lights off (to the right). Dp1 the day after CC and dp13 the day before the next CC. See Supplementary information (Fig S6) for corresponding data of the other 7 mice.

### Distance and speed of bouts of motion

Bouts of MOTS are local and usually cover a Euclidian distance of less than 2cm and motion is at low speed (Table 4). Locomotion usually covers 0.1 to 0.2 m, i.e., the distance from one side to the other of a cage and less than 1% of the locomotor bouts are longer than 1m (Table 4).

**Table 4.**

| Speed and distance made during bouts of movement |  |  |  |  |
| --- | --- | --- | --- | --- |
|  | Median | Mean | Range |  |
| Locomotor bout distance (m) | 0.139 | 0.174 | 0.013 | 3.726 |
| Locomotor bout velocity (m/s) | 0.026 | 0.028 | 0.002 | 0.076 |
| MOTS bout distance(m) | 0.006 | 0.015 | 0.000 | 0.810 |
| MOTS bout velocity (m/s) | 0.004 | 0.006 | 0.001 | 0.006 |

Distance made per day is about 250m during the dark hours and about 80m during daytime, except for the day of the cage change (Fig. 6A). The animal-to-animal variation in distance is larger during the dark vs. the light period of the day (*idem; and Suppo*rtive information (Fig. S7)). However, the average speed during locomotion is very similar (∼2.8 cm s^-1^) in darkness and day light (Fig. 6B).

**Fig 6.** (A) shows cumulative distance per 12 hrs during lights off (black line) and lights on (blue line), respectively. Values are average across the 10 animals during ZT 0-12 and ZT 12-24 each day with standard error indicated by bars. (B) shows corresponding data for average speed (±SEM) during lights on (blue) and lights off (black).

Locomotor bouts covering longer distance (>1m; Fig. 7A-F and Supportive information (Fig S8)) illustrates the variability of in-cage trajectories (Fig. 7 A-C) and the speed dynamics of locomotor bouts (Fig. 7 D-F). Speed ranges from a low of 0.01 m/s to 0.1m/s, and occasionally even higher speeds. Tracks also reveal occurrence of abnormal behaviours e.g., recursive locomotor activity (Fig. 7B, E).

**Fig 7.** A-C show longest recorded locomotor bout for mouse S1, S9 and S10, respectively during the observation period. In D-F, the corresponding speedograms are depicted. Track plots and speedograms by the trajar package in R. For tracks and speedogram of the other 7 mice see Supportive information (Fig S8).

### Spatial distribution of bouts’ starting point

The starting coordinates for rest and PA bouts were analysed to explore how the cage floor is used and to compare the day of the cage change (dp0) with the final day of the cage-change cycle (dp13)(Fig. 8 and Supportive information (Fig S9)). There is a considerable variability within and between mice, across a cage cycle and between cycles. Overall, there is a significant decrease by ∼40% (p<0.002; Table 5) in total number of bouts/day across the cage-change cycle, while the variation in bout reduction between cycles was not significant (Table 5).

**Table 5.**

| Reduction (%) of bouts dp13 vs dp0 |  |  |  |  |
| --- | --- | --- | --- | --- |
| Cage change cycle | Mean | SD | Median |  |
| Cycle 1 | -38.8 | 4.3 | -37.2 | n=10 |
| Cycle 2 | -44.0 | 3.5 | -43.5 | n=10 |
| Cycle 3 | -40.0 | 8.7 | -43.9 | n=10 |

Model: bout~cycle; F=2.1, p=0.14

**Fig 8.** Densitograms showing the distribution of bout starting coordinates during 24h on dp0 and dp13 in cage change cycles 1 – 3 (columns) for the S1, S7 and S8 mouse, respectively (rows). Cage front and rear and left (L) and right (R) have been indicated. Key to colour: Blue is long rest bout, orange is short rest bout, green is MOTS, and red is locomotor bout. Please see Supplementary information for corresponding metrics of the other mice kept in isolation.

In some cases (Fig. 8 and Supportive information (Fig S9)) there is an even distribution across the cage floor of the starting points on both the day of the cage-change and at the end of the cage change cycle although number of bouts and their relative abundance changes (see also below). In other cases, bout initiation tended to be more clustered, and this changed during as well as between cycles of the same mouse (e.g., case S7 in Fig. 8).

There was no clear preference for initiating bouts in the frontal vs rear area of the cage floor (Fig. 9 A-C; for definition of the cage floor sub fields see Fig. 1), the distribution is close to even except for the few long rest bouts which tended to be more frequent in the rear (Fig 9B). In contrast, bout initiation was infrequent in the central area of the cage floor (∼25%; Fig 9 D-F), regardless of the bout being rest or a movement.

**Fig 9.** A Boxplot showing the preference for bout initiation in the frontal field of the cage floor on dp0 and dp13 through cage change cycles 1-3. In B and C, the relative frequency of long rest and locomotor bouts, respectively, starting in the frontal field of the cage floor have been indicated. D-F show the corresponding boxplots when the cage floor was divided into a central and peripheral field of equal size (see also Fig. 1).

### Rhythmicity of rest and locomotion of single-housed mice

Mice follow a circadian rhythm of rest and PA, entrained to lights on and lights off in the laboratory environment (see Supportive information (Fig S1)). There is nocturnal peak of PA while daytime holds the highest density of long periods of rest (Fig. 10 A, C). This pattern of rest and PA reproduces closely across cages (Fig. 10 A, C) and between metric used (tracking, Fig. 10A; EAD, Fig. 10B) across the LD cycle.

**Fig 10.** A shows fraction of daily distance (m) per hours during the LD cycle for each mouse (thin coloured lines) and the average across the ten cages (thick black line). B shows the EAD per unit time when the animals are active during the LD cycle. Individual mice indicated by thin coloured lines, average across the group is the thick black line. C show the average fraction of each hour the mice spend in long rest (blue), short rest (orange), MOTS (grey) and in locomotion (yellow) across the LD cycle. Abscissa is Zeitgeber time with lights on 0-12 and lights off 12-24, the shift on to off is marked by a red vertical line.

Decomposing the datafiles into bouts of long and short rest, MOTS and locomotion provide the basis for the observed alteration in travelled distance (or EAD) and EAD across the circadian cycle (compare Fig. 10 A-B with C).

### Activity and rest by electrode activation density (EAD) in single and group housed mice

Consistent with the tracking data reported above, the EAD metric show that 75% of the time spent resting (80% of total file time) among single housed mice are bouts of long rest (≥40s; 59% of total time) having an average duration of ∼300s (Fig 11 A-C). The discrepancy between the tracking metric and the EAD appears to be due to EAD occurring during bouts when the mouse centroid coordinates does not change (c.f. Tables 2).

**Fig 11.** A-B Boxplots of fraction of total file spent at rest (no electrode activation) (A) and long rest (B) for the cohorts of female and male mice housed at different density (Fx1, Mx2, Fx3 etc). C-D show boxplots of duration and density of long rest bouts assessed by the EAD metric. As in A-B, female and male cohorts housed at different densities have been indicated.

As expected, increasing housing density decreases the amount of time spent in synchronized long rest (model: long rest ∼density * sex; whole model F=239, p<0.0001; and p<0.0001 for density; sex was not a significant factor p=0.514), mainly by reducing bout duration (Fig. 11C, F=130, p<0.0001; effect by density p<0.0001 and by sex, p<0.0001) but also the frequency (Fig. 11D, F=69, p<0.0001; effect by density p<0.0001 and sex p=0.0001). Eighty-two percent of the EAD occurred during bouts of motion, with 2/3^rd^ in locomotor bouts (Table 2). In Fig 12 the impact of housing density and sex on EAD is shown and the effect size (Hedge’s q) has been tabulated in Table 6. The stepwise increase in EAD is large (effect size∼2) and without any apparent difference between sexes (Table 6) until the density reaches four per cage. The effect size on EAD by increasing density from x3 female or x3 male mice up to x4 male mice was 1.4, which is only half of the effect size (∼3) recorded for the step up from x3 to x4 female mice (Table 6).

**Table 6.**
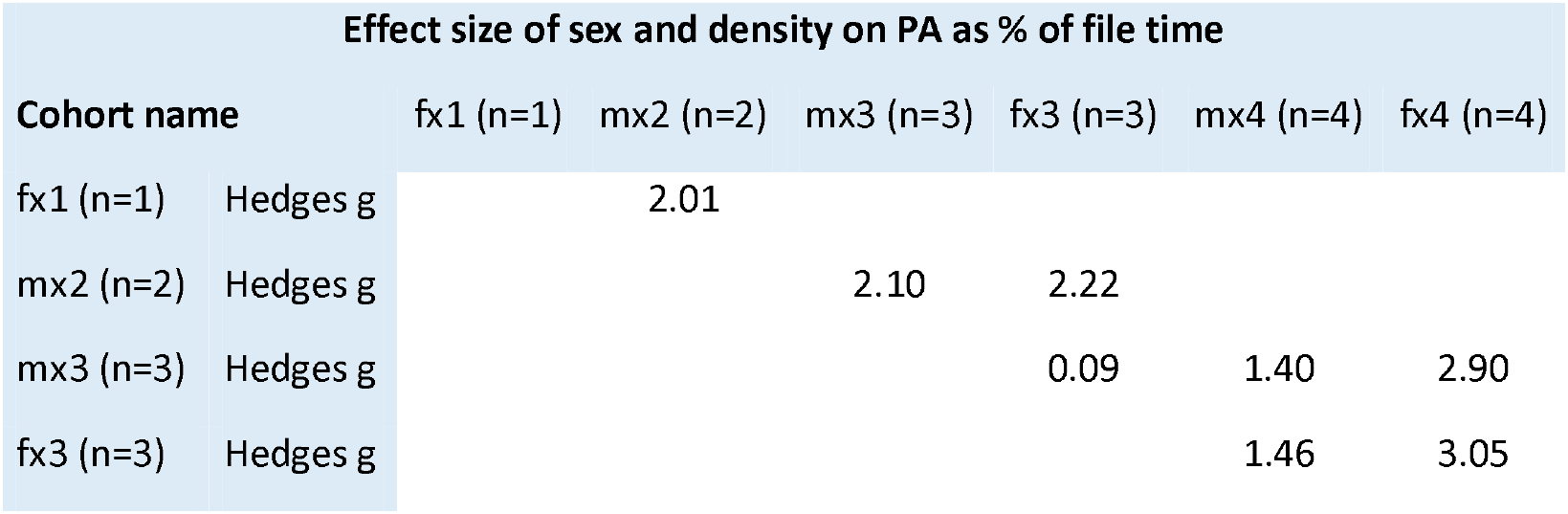
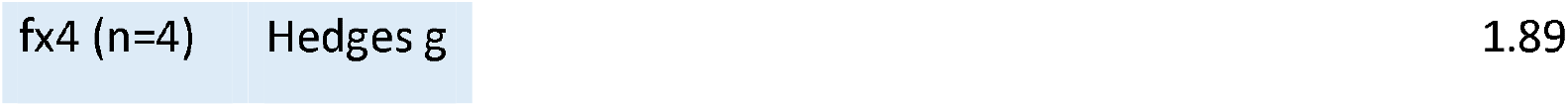

**Fig 12.** A-B Cumulative plots of time in PA assessed by EAD and the number of electrode activations observed (abscissa) as fraction of file time (ordinate) for each cohort of female (A) and male (B) cages (starting point of curve represent all bouts of rest (R). Housing density (x1, x2, x3, and x4) has been colour coded (key in panel A-B). Solid line represents cohort average value across cages and weeks of recording. The shaded area with the same colour indicates the standard deviation. Interrupted vertical lines indicate cut point values for rest (R) and fold change in electrode activations s-1.

### Rest and activity (EAD) across the LD and the cage-change cycle

As previously reported the amount of PA and rest per hour varies systematically across the LD cycle (Fig. 13) and the proportion of synchronized-rest decreases as housing density increases. Above we showed that bouts of rest and PA were impacted by the cage change (Figs 4 and 6) affecting both distance moved and bout composition. When we use EAD as metric (Figs 12, Table 6) we obtained corresponding results for the single housed mice and a stepwise increase of effect size as we increase housing density up to four. In line with the results using aggregated EAD data (Fig. 12 A-B and Table 6), there was a significant difference between sexes at density x4 but not at x3 with respect to both time spent in PA and synchronized rest (Fig. 14 A-B). In all the different housing densities studied here, there was a marked impact by the cage change in both sexes on time spent in PA and synchronized-rest, respectively (*idem; see also* Supportive information (Fig S10) for effect size).

**Fig 13.** Panels show the average fraction of each hour spent in PA (yellow) and at rest (blue) across the LD cycle for male and female mice housed at different densities. Rest i.e., no electrode activation and PA when electrode activations occur. Abscissa is Zeitgeber time with lights on 0-12 and lights off 12-24, the shift on to off is marked by a red vertical line.

**Fig 14.** A show fraction of file time spent in PA (≥ electrodes activated s-1) across cages per day (dp) of the cages change cycles 1-3 for male and female mice housed at different densities (x1: n=1, x2 n=2, x3 n=3 and x4 n=4). Comparison of female and male mice at density n=3 and n=4, respectively, revealed a significant difference at density n=4 but not when density =3 (model: Time in PA ∼ sex * dp * CC; density =4 F=36.6; n=3 F=0.07). See also Supplementary information (Fig S10) for plot of relative effect size of dp and CC across. B show fraction of file time spent in long rest bouts (<1 electrodes activated s-1; ≥40 s duration) across cages and cages-change cycles (CC 1-3) per day (dp0-dp13) for male and female mice housed at different densities (x1 to x4). Comparison of female and male mice at density n=3 and n=4, respectively, revealed a significant difference in synchronized long rest bout time at density n=4 but not when n=3 (model: Time in long rest ∼ sex * dp * CC; n=4 F=7.6; n=3 F=0.79). See Supportive information for plot of relative effect size of dp and CC (Fig. S10). C show average number of unique electrodes activated s-1 across cages per day of the cages change cycles 1-3 for male (blue) and female (red) mice housed at different densities (x1 to x4). Comparison of female and male mice (model: No unique electrodes ∼ sex * dp * CC) revealed a significant difference at density =4 (F=41, p=1.9E-5) but not at x3 (F=1; p=0.33). See Supportive information for plot of relative effect size of dp and CC (Fig. S10).

EAD increases (Fig. 14A, see also Supportive information (Fig S3)) with housing density and in parallel a larger number of unique electrodes are activated during a PA bout (Fig 14C), suggesting that number of unique electrodes involved in a PA bout covariate with EAD.

With a housing density of four female or male mice, the relative effect size of days post cage change on PA and rest across the cage change cycle displayed a biphasic trajectory with a second distinct infliction on the day when neighbouring cages were subjected to a cage change. Such inflictions were occasionally also evident in cages with three animals but not a consistent feature through cycles (*idem) and, furt*hermore, not seen with lower housing densities (Fig. 15).

**Fig 15.** The panels left-to-right show relative effect size (RTE, ordinate) on average daily EAD by housing density (n), days post cage change (abscissa; dp0-13) and cage-change cycle (CC 1-3; colour coded black, red, and green, in each panel) within sexes (group of panels). Whole model: EAD ∼ N * dp * CC; for females F=163.7, p=7.6E-15; for males F=28, p=3.3E-7. In both sexes, density (n) had the strongest impact on EAD, followed by dp, while impact by cage change cycle was only significant in females. NB: In males only two cycles could be compared across densities. Note that at N=x4 EAD show a biphasic trajectory across all cycles in both sexes (red arrows, for further information see

EAD occurrence in the frontal vs the rear part of the cage floor was rather similar with 5%-10% more EADs in the rear of the single-housed mice. In two out of three cycles, the percentage of EADs in the frontal area increased by 5-10% towards the end of the cage change cycle (Table 7).

**Table 7.**

|  | Frontality of EAD during and across cage change cycle(s) |  |  |  |  |  |  |  |
| --- | --- | --- | --- | --- | --- | --- | --- | --- |
| | dp0 | | | dp13 | | | $\Delta$ % dp0-dp13 | |
|  | Mean | Median | SD | Mean | Median | SD | Mean | n |
| Cycle 1 | 41.7 | 38.4 | 11.3 | 48.0 | 48.6 | 8.1 | 6.3 | 10 |
| Cycle2 | 44.0 | 41.6 | 9.8 | 52.5 | 50.6 | 10.4 | 8.5 | 10 |
| Cycle3 | 47.9 | 49.9 | 8.2 | 45.6 | 45.7 | 19.4 | -2.3 | 10 |

## Discussion

### General comments

In this study we use decomposition analysis of longitudinal data recorded with a DVC HCM system covering six weeks to assess pattern of rest and PA of single and group housed C57BL/6J mice. The decomposition of the record files was based exclusively on the occurrence or absence of a change between samples of the mouse centroid position (≥1 mm). Further subdivision of bouts were based on bout duration (short and long rest, split at 40s) and with PA bouts into MOTS (local movement) and locomotion (the bouts’ radial distance was larger, or equal to, one average stride length for this strain and sex)[17, 19]. Thus, the decomposition into bouts did not compromise the resolution of the system. In single housed mice, we extracted both track and EAD metrics from the raw data to assess degree of covariation. As confirmed here (see [16]), EADs and distance by tracking per hour correlate closely but there is not a perfect match (∼4% of file time differed) and, further, in bouts of movements the relation between EADs and made distance was different between PA bout types because MOTS of longer duration associated with high EAD despite very short distance made. With tracking it is straight forward to differentiate between MOTS and locomotion, however, associating EAD with number of different electrodes activated and the temporal succession of electrode activations during a bout, may prove useful as guidance to separate locomotor and MOTS bouts with the EAD metric. As with tracking, however, this can only be applied when animals are single-housed [15, 16]. In the discordant segments of the aligned [EAD and tracking] files, electrode activations were recorded during periods when the tracking did not recognize any movement (<1mm, ∼resolution of the system [15]), i.e. during bouts of rest. Although this deviation between records is only a few percent of the total file time, the metrics on time spent in long rest differed (EAD: 59% and tracking: 67% of recorded time) while the difference of total time in rest was only ∼1%. A plausible explanation is that EAD is a more sensitive metric of PA, possibly responding to changes in body posture and movements of the tail. Such electrode activations may not be sufficient to alter the x, y position of the mouse centroid. This led us to interpret long rest bouts as sleep only if both metrics indicated that the animal did not move (see also [21] on criteria stringency of being still). Thus, the time budget for rest of single housed (C57Bl/6) mice was ∼80% considering the two metrics extracted, and sleep (NREM+REM) 59% (based on EAD).

With the definition for PA bout types used here, about equal time is used for MOTS and locomotion. With DVC, the tracking metric is more informative than the EAD but we recommend to extract both and to use them in parallel (as we did here to define rest as sleep). EAD can also be used when animals are co-housed to relay information about bouts of synchronized-rest and group PA (density and spatial distribution of activation) of the animals (see also below) and have been used to monitor circadian rhythm, impact by husbandry routines, and progression of biological processes and diseases[12, 27, 29, 33, 42, 43].

### Bouts of rest and PA

Single housed mice generate ∼6500 bouts of rest and PA per day, this is in agreement with data for C57BL/6 obtained by a HCM system using force plates[44]. Eighty percent are bouts of PA corresponding to only ∼20% of the file time (see also [45, 46], while only 2.5% of the bouts are long rest bouts considered sleep episodes making up 59% of the file time. Sleep is divided into non-REM sleep (NREM; in mice >90%) and REM sleep (REM; <10%). The gold standard to decide on state of vigilance is to record gross cortical brain activity by surface electrodes (EEG), eye-movements, and muscle tone (EMG). Pack et al [47] found by comparing EEG and EMG recording with CCD a >90% agreement between long periods of rest (CCD; ≥30s) and EEG-EMG pattern of NREM and REM sleep. These observations were later confirmed and extended by correlative assessment of home-cage periods of long rest using piezo-electric sensor[48], IR-sensor[20, 49], CCD[21, 22], and electric-field sensor [50] based HCM techniques combined with EEG-EMG recordings. The results of these studies show a correlation >0.9 between NREM and REM sleep, on the hand, and the animals being still ≥40s, on the other. This behavioural criterion for sleep has subsequently been used in a number of studies[51-55]. We conclude based on our results and those obtained with other HCM techniques (*idem) that bout*s of inferred sleep (no distinction made here between NREM and REM) in single housed mice of this strain commonly have a duration of 300-400s but can exceed 1000s and occur with an average density of ∼120 bouts per day preferentially during lights on. Moreover, our bout actigrams revealed that sleep bouts were clustered at 4-6 time periods during lights on, and 1-2 episodes during lights off. Interventions like a cage change induces considerable fragmentation of bouts, affecting those considered to be sleep (see also [28, 56-58]).

With co-housing the time spent in synchronized long rest (no EAD) decrease, in pair housing to 45-50% of file time, with trios down to ∼35% and no difference between sexes. Still at a housing density of four mice per cage ≥25% of the file time is spent in synchronized long rest, inferred to be sleep. At this density, males had significantly shorter synchronized long rest than females.

As highlighted by Golani and co-workers [17], bouts of physical activity in mice can be divided into locomotion and local movements (MOTS; see also [44, 45]). Local movements comprise a range of behavioural entities not possible to decode with the DVC system. These entities include feeding, drinking, rearing, and grooming (*idem and [46])*. MOTS make up ∼75% of the PA bouts and half of the file time devoted to PA. They are more prevalent during lights off (∼65% of all daily MOTS) and in agreement with data recorded with other types of HCM systems[44, 45] they usually have a short duration (≤2s), and cover typically a short distance (<2cm) at low speed (<0.6 cm s^-1^). Our data shows that a fraction of the MOTS have a longer duration, and that these bouts associate with high values for EAD. Such bouts may correspond to e.g., eating bouts. MOTS bouts increase in response to a cage change and in the responses to lights on/off during the LD cycle, and cover a cumulative distance of 40-70m per day, i.e. ∼15-20% of the daily moved distance (see also[44]).

We recorded 1000-2000 bouts of locomotion per day in single housed mice. As expected, few occurs during lights on because mice are nocturnal. The typical locomotor bout covers a distance of 0.1-0.2 m with a speed of ∼3 cm s^-1^, which agrees with previously published data recorded with DVC and other types of HCM systems[53, 56, 59, 60] but lower than those reported by [44, 45]. This discrepancy in average speed may relate to differences in the definition of a locomotor bout (speed or distance). As shown here, speed varies (range∼0.01-0.1 m s^-1^) during a locomotor bout, and the trajectory and bout speedogram will unmask abnormal recursive motor behaviours (fits of stereotypy). Distance travelled per hour is in the range from <1m h^-1^ (day time during periods of rest) up to 40m h (in response to lights on, and cage change), decays across days of the cage change cycle and show considerable differences (one fold) between [single housed] mice. The daily average distance covered by the mice was ∼330m in this study, which is within the range (∼150-750 m 24h) of previously published data [16, 44, 45, 56, 59, 60]. The difference noted in average speed between our results, including that we did not find a difference in speed during movements day time vs night time, and those previously published using DVC is due to that we used decomposition into bouts, while Iannello’s [16] data is average across all movements and rest per unit time, and Shenk et al.[60] used a different definition of locomotion.

### Rest and PA across the LD cycle, and the use of the cage floor

In mammals, the pattern of PA and rest follows different rhytmicities (for references see [29]). Universal is the circadian rhythm which entrains to Earth’s Day and Night (*idem). HCM syst*ems are ideal for the purpose of analysing behavioural rhytmicities over extended periods of time in small rodents [29, 33] and may prove to be a good substitute to the current gold standard of using running wheels in studies of the circadian rhythm[61]. Mice are nocturnal and will rest during day time (lights on) while they are active and feed during night time (lights off). Our data show that across the LD cycle, the driving force is the clustering of long rest bouts to day time, while all other bout types increase in prevalence during night time. We also confirm that the responses to lights on/off appears insensitive to if lightning change suddenly or through a dawn and dusk transition period[27].

Although less informative than tracking EAD revealed the same pattern of rest and PA and, further, that the amount of time spent in PA bouts increase stepwise up to a density of four mice without any apparent difference between sexes [for the C57BL/6J strain of mice]. We confirm previous observations that mice housed 4 to a cage, the level of PA and rest differ between sexes[12, 27].

In a recent publication we showed that male mice housed 4 to a cage tend to constrain the use of the cage floor area across the cage change cycle and that this may be related to the location of the in-cage latrine(s)[27]. Females at the same housing density did not show the same degree of spatial clustering of PA and, importantly, this was not evident when males were housed in pairs (*idem). Single h*oused mice, tend to locate long rest into the rear half of the cage (mainly during lights on), while all other bout types (97.5%) were spread across the front and rear of the cage floor but with a significant portion of them (75%) originating (and terminating) in the peripheral part (50% of floor area) of the cage floor. Thus, housing C57BL/6J mice at densities ≥4 per cage seems to alter in cage behaviour and induce sex differences, combined this may indicate a crowdedness stress that should be taken into account when designing experiments and comparing study results.

### Decomposition of longitudinal HCM records into bouts of rest and PA may have significant value in translational research

The introduction of small wearable devices (accelerometers, more recently smart watches and similar items) that can be carried on the wrist, thigh, or trunk without disrupting normal activity has made it possible to collect large amounts of longitudinal data on rest and activity from healthy and sick, growing and aging humans, to extract metrics useful as objective biomarkers of development and ageing, disease progression, outcome prediction as well as monitoring impact by intervention [34, 62-75]. So far, only few accelerometer studies on humans encompasses variation in activity and rest bouts across the circadian cycle[34, 63, 67, 68, 72]. However, as smart watches and similar devices are likely to replace the older more unpractical accelerometers this will likely change. The number of papers by year presenting accelerometer data in humans has increased x25 from year 2001 to 2020 (source PubMed). Work is ongoing to develop recommendations to standardize accelerometer records as well as new tools by which the data can be analysed [31, 32, 63, 76-82]. Similar initiatives are currently ongoing in the realm of HCM of animal models in the life sciences with a recently started COST action in EU (TeaTime) and the North American 3Rs collaborative (Na3RsC).

The most common approach so-far to analyse accelerometer/smartwatch raw data is by decomposition analysis using cut-point values to stratify the data into bouts of sleep (SL) and sedentary behaviour (SB), low-medium and vigorous physical activity (PA) or different combinations of these categories (*idem). The freq*uency and duration as well as accumulation pattern of different bout types and the composition of bouts are then compared over time and/or between groups. With GPS tracking becoming a more frequent feature of wearable devices also distance made and speedograms will be possible to retrieve as indices. Similarly, data generated by a variety of HCM systems have recently been used in efforts to identify behavioural indices of ageing[29, 83], impact of disease progression[42, 43, 60, 84, 85], genetic modification[44, 48, 51, 53, 86], and insults[59, 60, 87].

Since the data generated by wearable devices in humans and by non-intrusive scalable HCM systems in animals essentially overlap, it should be feasible to agree on sets of metrics that will serve as equivalent biomarkers for different conditions and biological process in both humans and small rodent models used to study human conditions.

### Limitations of this study

With few exceptions[9, 88, 89], scalable HCM systems such as the DVC used here can provide detailed metrics only when animals are kept in isolation (for references see Introduction and above). Mice are normally living in groups and as stated in the EU Directive63/2010 there are ethical reasons to avoid single housing of laboratory rodents, since evidence indicate a depreciation of animal welfare and that isolation may alter animals’ mental capacity and spontaneous behaviour [90-97]. However, some studies especially on male mice have questioned this and showed that the welfare or behaviour must not always be depreciated and, furthermore, depends on the context [95, 98-102].

To enable assessment of data extracted from the raw electrode output of the DVC, we used a cohort of single housed females as the main subjects of this study. Our results indicate that with co-housing, PA (EAD) increased in a stepwise fashion from single housing to pair housing and from pairs up to trios, regardless of sex for this strain of mice. Thus, our data suggests that welfare depreciation experienced due to single housing did not seriously affect the mice’s daily amount of PA to a significant degree. It remains roughly proportional to number of animals in the cage up to a density of four animals.

Both the spatial (12 electrodes spaced apart) and temporal resolution (4 Hz) of the system are low compared to video-based solutions (usually ≥ 15 FPS and HDMI) but has the advantage of low demands on IT infrastructure, data storage and processing capacity. Except for the rather few long bouts of rest, the different bout types had a frequency in the range of 0.05-1 Hz. Thus, the over-sampling at 4 Hz should be sufficient to delineate the bouts accurately. The HCM system is fully automatable, scalable, and non-intrusive. Similar, to RFID (Intellicage, TSE; e.g. [4]), IR-beam (Actimot, TSE; e.g.[8]) and Force plate (Metris, Labora; e.g. [6])-based systems, the output does not have high dimensionality. Thus, this type of system cannot capture behavioural trait details, only basic metrics as animal localisation, rest, and PA. An additional limitation is that recording of PA (and rest) with the DVC system is restricted to the floor of the cage and activities such as e.g. climbing is not recognized. Moreover, it cannot be excluded that data of PA was not picked-up when the animal was on top of shuffled piles of bedding and enrichment materials.

Still, as shown here tracking and EADs can be quite informative when the animals are kept in isolation, while housed in groups the information provided (by EAD metric) apply only to the group not individual animals.

### Concluding remarks

We show that data on a variety of parameters such as sleep pattern, locomotor activity, bout fragmentation, spatial distribution of rest and PA, the circadian rhythm and changes to these metrics induced by interventions, can be extracted in near real-time from scalable HCM systems like the DVC using standard desk top computers. Longitudinal data can easily be generated and retrieved on a large scale serving both the care and welfare of the experimental animals, and the research conducted on them. The output from these systems compares well with data generated by wearable devices on humans and may, thus, form a basis for translatable behavioural biomarkers.

## Acknowledgements

Help and advice from technical staff at the Wallenberg laboratory facility at Karolinska Institutet (KI) and the ECF facility at Stockholm University (SU) are gratefully acknowledged. We thank Mara Rigamonti and Giorgio Rosati of the DVC division at Tecniplast SpA, Italy, for technical support and discussions, and the assistance with transfer of the raw data files with millisecond resolution.

## Cited literature

1. Todd JT. A selective look at some pre-skinnerian cumulative recording systems and cumulative records in physiology and psychology. Mexican Journal of Behavior Analysis. 2017;43(2):137–63.

2. Robinson SF, Pauly JR, Marks MJ, Collins AC. An analysis of response to nicotine infusion using an automated radiotelemetry system. Psychopharmacology (Berl). 1994;115(1-2):115–20. doi: 10.1007/bf02244760. PubMed PMID: 7862882.

3. Tamborini P, Sigg H, Zbinden G. Quantitative analysis of rat activity in the home cage by infrared monitoring. Application to the acute toxicity testing of acetanilide and phenylmercuric acetate. Arch Toxicol. 1989;63(2):85–96. doi: 10.1007/bf00316429. PubMed PMID: 2730344.

4. Dell’omo G, Shore RF, Lipp HP. An automated system, based on microchips, for monitoring individual activity in wild small mammals. The Journal of experimental zoology. 1998;280(1):97–9. PubMed PMID: 9437856.

5. Krackow S, Vannoni E, Codita A, Mohammed AH, Cirulli F, Branchi I, et al. Consistent behavioral phenotype differences between inbred mouse strains in the IntelliCage. Genes Brain Behav. 2010;9(7):722–31. Epub 2010/06/10. doi: 10.1111/j.1601-183X.2010.00606.x. PubMed PMID: 20528956.

6. Altun M, Bergman E, Edstrom E, Johnson H, Ulfhake B. Behavioral impairments of the aging rat. Physiol Behav. 2007;92(5):911–23. Epub 2007/08/07. doi: S0031-9384(07)00263-6 [pii] 10.1016/j.physbeh.2007.06.017. PubMed PMID: 17675121.

7. Grieco F, Bernstein BJ, Biemans B, Bikovski L, Burnett CJ, Cushman JD, et al. Measuring Behavior in the Home Cage: Study Design, Applications, Challenges, and Perspectives. Front Behav Neurosci. 2021;15:735387. Epub 20210924. doi: 10.3389/fnbeh.2021.735387. PubMed PMID: 34630052; PubMed Central PMCID: PMCPMC8498589.

8. Fahlstrom A, Zeberg H, Ulfhake B. Changes in behaviors of male C57BL/6J mice across adult life span and effects of dietary restriction. Age (Dordr). 2012;34(6):1435–52. Epub 2011/10/13. doi: 10.1007/s11357-011-9320-7. PubMed PMID: 21989972; PubMed Central PMCID: PMC3528371.

9. Redfern WS, Tse K, Grant C, Keerie A, Simpson DJ, Pedersen JC, et al. Automated recording of home cage activity and temperature of individual rats housed in social groups: The Rodent Big Brother project. PloS one. 2017;12(9):e0181068–e. doi: 10.1371/journal.pone.0181068. PubMed PMID: 28877172.

10. Richardson CA. The power of automated behavioural homecage technologies in characterizing disease progression in laboratory mice: A review. Applied Animal Behaviour Science. 2015;163:19–27. doi: 10.1016/j.applanim.2014.11.018.

11. Bains RS, Cater HL, Sillito RR, Chartsias A, Sneddon D, Concas D, et al. Analysis of individual mouse activity in group housed animals of different inbred strains using a novel automated home cage analysis system. Frontiers in Behavioral Neuroscience. 2016;10(JUN). doi: 10.3389/fnbeh.2016.00106.

12. Pernold K, Iannello F, Low BE, Rigamonti M, Rosati G, Scavizzi F, et al. Towards large scale automated cage monitoring - Diurnal rhythm and impact of interventions on in-cage activity of C57BL/6J mice recorded 24/7 with a non-disrupting capacitive-based technique. PLoS One. 2019;14(2):e0211063. Epub 2019/02/05. doi: 10.1371/journal.pone.0211063. PubMed PMID: 30716111; PubMed Central PMCID: PMCPMC6361443 Buguggiate (Va), Italy) is a commercial company selling the DVC system. However, this does not alter the authors’ adherence to all the PLOS ONE policies on sharing data and materials. We have read the journal’s policy and the authors of this manuscript have no competing interests.

13. Baran SW, Bratcher N, Dennis J, Gaburro S, Karlsson EM, Maguire S, et al. Emerging Role of Translational Digital Biomarkers Within Home Cage Monitoring Technologies in Preclinical Drug Discovery and Development. Front Behav Neurosci. 2021;15:758274. Epub 20220214. doi: 10.3389/fnbeh.2021.758274. PubMed PMID: 35242017; PubMed Central PMCID: PMCPMC8885444.

14. Gaburro S, Winter Y, Loos M, Kim JJ, Stiedl O. Editorial: Home Cage-Based Phenotyping in Rodents: Innovation, Standardization, Reproducibility and Translational Improvement. Front Neurosci. 2022;16:894193. Epub 20220623. doi: 10.3389/fnins.2022.894193. PubMed PMID: 35812217; PubMed Central PMCID: PMCPMC9261870.

15. Klein C, Budiman T, Homberg JR, Verma D, Keijer J, van Schothorst EM. Measuring Locomotor Activity and Behavioral Aspects of Rodents Living in the Home-Cage. Front Behav Neurosci. 2022;16:877323. Epub 20220407. doi: 10.3389/fnbeh.2022.877323. PubMed PMID: 35464142; PubMed Central PMCID: PMCPMC9021872.

16. Iannello F. Non-intrusive high throughput automated data collection from the home cage. Heliyon. 2019;5(4):e01454. Epub 2019/04/19. doi: 10.1016/j.heliyon.2019.e01454. PubMed PMID: 30997429; PubMed Central PMCID: PMCPMC6451168.

17. Drai D, Kafkafi N, Benjamini Y, Elmer G, Golani I. Rats and mice share common ethologically relevant parameters of exploratory behavior. Behav Brain Res. 2001;125(1-2):133–40. Epub 2001/10/30. doi: 10.1016/s0166-4328(01)00290-x. PubMed PMID: 11682104.

18. Drai D, Golani I. SEE: a tool for the visualization and analysis of rodent exploratory behavior. Neurosci Biobehav Rev. 2001;25(5):409–26. Epub 2001/09/22. doi: 10.1016/s0149-7634(01)00022-7. PubMed PMID: 11566479.

19. Fahlstrom A, Yu Q, Ulfhake B. Behavioral changes in aging female C57BL/6 mice. Neurobiol Aging. 2009. Epub 2009/12/17. doi: S0197-4580(09)00362-5 [pii] 10.1016/j.neurobiolaging.2009.11.003. PubMed PMID: 20005598.

20. Angelakos CC, Watson AJ, O’Brien WT, Krainock KS, Nickl-Jockschat T, Abel T. Hyperactivity and male-specific sleep deficits in the 16p11.2 deletion mouse model of autism. Autism Res. 2017;10(4):572–84. Epub 20161014. doi: 10.1002/aur.1707. PubMed PMID: 27739237; PubMed Central PMCID: PMCPMC6201314.

21. Fisher SP, Godinho SI, Pothecary CA, Hankins MW, Foster RG, Peirson SN. Rapid assessment of sleep-wake behavior in mice. J Biol Rhythms. 2012;27(1):48–58. Epub 2012/02/07. doi: 10.1177/0748730411431550. PubMed PMID: 22306973; PubMed Central PMCID: PMCPMC4650254.

22. McShane BB, Galante RJ, Biber M, Jensen ST, Wyner AJ, Pack AI. Assessing REM Sleep in Mice Using Video Data. Sleep. 2012;35(3):433–42. doi: 10.5665/sleep.1712.

23. Brown LA, Hasan S, Foster RG, Peirson SN. COMPASS: Continuous Open Mouse Phenotyping of Activity and Sleep Status. Wellcome open research. 2016;1:2-. doi: 10.12688/wellcomeopenres.9892.2. PubMed PMID: 27976750.

24. Tang X, Xiao J, Parris BS, Fang J, Sanford LD. Differential effects of two types of environmental novelty on activity and sleep in BALB/cJ and C57BL/6J mice. Physiol Behav. 2005;85(4):419–29. Epub 2005/07/16. doi: 10.1016/j.physbeh.2005.05.008. PubMed PMID: 16019041.

25. Erratum to “FELASA recommendations for the health monitoring of mouse, rat, hamster, guinea pig and rabbit colonies in breeding and experimental units”. Lab Anim. 2015;49(1):88. Epub 2014/09/04. doi: 10.1177/0023677214550970. PubMed PMID: 25181995.

26. Mahler Convenor M, Berard M, Feinstein R, Gallagher A, Illgen-Wilcke B, Pritchett-Corning K, et al. FELASA recommendations for the health monitoring of mouse, rat, hamster, guinea pig and rabbit colonies in breeding and experimental units. Lab Anim. 2014;48(3):178–92. Epub 2014/02/06. doi: 10.1177/0023677213516312. PubMed PMID: 24496575.

27. Ulfhake B, Lerat H, Honetschlager J, Pernold K, Rynekrová M, Escot K, et al. A multicentre study on spontaneous in-cage activity and micro-environmental conditions of IVC housed C57BL/6J mice during consecutive cycles of bi-weekly cage-change. PLoS One. 2022;17(5):e0267281. Epub 20220525. doi: 10.1371/journal.pone.0267281. PubMed PMID: 35613182; PubMed Central PMCID: PMCPMC9132304.

28. Saré RM, Lemons A, Torossian A, Beebe Smith C. Noninvasive, High-throughput Determination of Sleep Duration in Rodents. J Vis Exp. 2018;(134). Epub 20180418. doi: 10.3791/57420. PubMed PMID: 29733321; PubMed Central PMCID: PMCPMC6100687.

29. Pernold K, Rullman E, Ulfhake B. Major oscillations in spontaneous home-cage activity in C57BL/6 mice housed under constant conditions. Sci Rep. 2021;11(1):4961. Epub 2021/03/04. doi: 10.1038/s41598-021-84141-9. PubMed PMID: 33654141; PubMed Central PMCID: PMCPMC7925671.

30. Zucker I. Circannual rhythms Mammals. Circannual rhythms Mammals in Handbook of Behavioral Neurobiology] Handbook of Behavioral Neurobiology Vol 122001. p. 509–29.

31. Leroux A, Di J, Smirnova E, McGuffey EJ, Cao Q, Bayatmokhtari E, et al. Organizing and analyzing the activity data in NHANES. Stat Biosci. 2019;11(2):262–87. Epub 2020/02/13. doi: 10.1007/s12561-018-09229-9. PubMed PMID: 32047572; PubMed Central PMCID: PMCPMC7012355.

32. Di J, Spira A, Bai J, Urbanek J, Leroux A, Wu M, et al. Joint and Individual Representation of Domains of Physical Activity, Sleep, and Circadian Rhythmicity. Stat Biosci. 2019;11(2):371–402. Epub 2020/05/23. doi: 10.1007/s12561-019-09236-4. PubMed PMID: 32440309; PubMed Central PMCID: PMCPMC7241438.

33. Bains RS, Wells S, Sillito RR, Armstrong JD, Cater HL, Banks G, et al. Assessing mouse behaviour throughout the light/dark cycle using automated in-cage analysis tools. J Neurosci Methods. 2018;300:37–47. Epub 20170426. doi: 10.1016/j.jneumeth.2017.04.014. PubMed PMID: 28456660; PubMed Central PMCID: PMCPMC5909039.

34. Xiao L, Huang L, Schrack JA, Ferrucci L, Zipunnikov V, Crainiceanu CM. Quantifying the lifetime circadian rhythm of physical activity: a covariate-dependent functional approach. Biostatistics. 2015;16(2):352–67. Epub 2014/11/02. doi: 10.1093/biostatistics/kxu045. PubMed PMID: 25361695; PubMed Central PMCID: PMCPMC4804116.

35. Nakagawa S, Cuthill IC. Effect size, confidence interval and statistical significance: a practical guide for biologists. Biol Rev Camb Philos Soc. 2007;82(4):591–605. Epub 2007/10/20. doi: 10.1111/j.1469-185X.2007.00027.x. PubMed PMID: 17944619.

36. Erceg-Hurn DM, Mirosevich VM. Modern robust statistical methods: an easy way to maximize the accuracy and power of your research. Am Psychol. 2008;63(7):591–601. doi: 10.1037/0003-066X.63.7.591. PubMed PMID: 18855490.

37. Noguchi K, Gel, R. L., Brunner, E., Konietschke, F. nparLD: An R Software Package for the Nonparametric Analysis of Longitudinal Data in Factorial Experiments. Journal of Statistical Software. 2012;50(12):1–23.

38. Cliff N. Dominance statistics: Ordinal analyses to answer ordinal questions. Psychological Bulletin. 1993;114(3):494–509. doi: 10.1037/0033-2909.114.3.494.

39. McGraw KO, Wong SP. A common language effect size statistic. Psychological Bulletin. 1992;111(2):361–5. doi: 10.1037/0033-2909.111.2.361.

40. Vargha A, Delaney HD. A Critique and Improvement of the “CL” Common Language Effect Size Statistics of McGraw and Wong. Journal of Educational and Behavioral Statistics. 2000;25(2):101–32. doi: 10.2307/1165329.

41. Grissom RJ, Kim JJ. Review of assumptions and problems in the appropriate conceptualization of effect size. Psychol Methods. 2001;6(2):135–46. doi: 10.1037/1082-989x.6.2.135. PubMed PMID: 11411438.

42. Golini E, Rigamonti M, Iannello F, De Rosa C, Scavizzi F, Raspa M, et al. A Non-invasive Digital Biomarker for the Detection of Rest Disturbances in the SOD1G93A Mouse Model of ALS. Front Neurosci. 2020;14:896. Epub 2020/09/29. doi: 10.3389/fnins.2020.00896. PubMed PMID: 32982678; PubMed Central PMCID: PMCPMC7490341.

43. Vagima Y, Grauer E, Politi B, Maimon S, Yitzhak E, Melamed S, et al. Group activity of mice in communal home cage used as an indicator of disease progression and rate of recovery: Effects of LPS and influenza virus. Life Sciences. 2020;258:118214. doi: https://doi.org/10.1016/j.lfs.2020.118214.

44. Chaudoin TR, Bonasera SJ. Mice lacking galectin-3 (Lgals3) function have decreased home cage movement. BMC Neurosci. 2018;19(1):27. Epub 2018/05/03. doi: 10.1186/s12868-018-0428-x. PubMed PMID: 29716523; PubMed Central PMCID: PMCPMC5930520.

45. Knibbe-Hollinger JS, Fields NR, Chaudoin TR, Epstein AA, Makarov E, Akhter SP, et al. Influence of age, irradiation and humanization on NSG mouse phenotypes. Biol Open. 2015;4(10):1243–52. Epub 20150909. doi: 10.1242/bio.013201. PubMed PMID: 26353862; PubMed Central PMCID: PMCPMC4610222.

46. Goulding EH, Schenk AK, Juneja P, MacKay AW, Wade JM, Tecott LH. A robust automated system elucidates mouse home cage behavioral structure. Proc Natl Acad Sci U S A. 2008;105(52):20575–82. Epub 20081223. doi: 10.1073/pnas.0809053106. PubMed PMID: 19106295; PubMed Central PMCID: PMCPMC2634928.

47. Pack AI, Galante RJ, Maislin G, Cater J, Metaxas D, Lu S, et al. Novel method for high-throughput phenotyping of sleep in mice. Physiol Genomics. 2007;28(2):232–8. Epub 20060919. doi: 10.1152/physiolgenomics.00139.2006. PubMed PMID: 16985007.

48. Lord JS, Gay SM, Harper KM, Nikolova VD, Smith KM, Moy SS, et al. Early life sleep disruption potentiates lasting sex-specific changes in behavior in genetically vulnerable Shank3 heterozygous autism model mice. Mol Autism. 2022;13(1):35. Epub 20220829. doi: 10.1186/s13229-022-00514-5. PubMed PMID: 36038911; PubMed Central PMCID: PMCPMC9425965.

49. Brown LA, Banks GT, Horner N, Wilcox SL, Nolan PM, Peirson SN. Simultaneous Assessment of Circadian Rhythms and Sleep in Mice Using Passive Infrared Sensors: A User’s Guide. Curr Protoc Mouse Biol. 2020;10(3):e81. doi: 10.1002/cpmo.81. PubMed PMID: 32865891.

50. Kloefkorn H, Aiani LM, Lakhani A, Nagesh S, Moss A, Goolsby W, et al. Noninvasive three-state sleep-wake staging in mice using electric field sensors. J Neurosci Methods. 2020;344:108834. Epub 2020/07/04. doi: 10.1016/j.jneumeth.2020.108834. PubMed PMID: 32619585; PubMed Central PMCID: PMCPMC7454007.

51. Saré RM, Harkless L, Levine M, Torossian A, Sheeler CA, Smith CB. Deficient Sleep in Mouse Models of Fragile X Syndrome. Front Mol Neurosci. 2017;10:280. Epub 20170901. doi: 10.3389/fnmol.2017.00280. PubMed PMID: 28919851; PubMed Central PMCID: PMCPMC5585179.52.

52. Saré RM, Lemons A, Song A, Smith CB. Sleep Duration in Mouse Models of Neurodevelopmental Disorders. Brain Sci. 2020;11(1). Epub 20201230. doi: 10.3390/brainsci11010031. PubMed PMID: 33396736; PubMed Central PMCID: PMCPMC7824512.53.

53. Schirmer C, Abboud MA, Lee SC, Bass JS, Mazumder AG, Kamen JL, et al. Home-cage behavior in the Stargazer mutant mouse. Sci Rep. 2022;12(1):12801. Epub 20220727. doi: 10.1038/s41598-022-17015-3. PubMed PMID: 35896608; PubMed Central PMCID: PMCPMC9329369.

54. Siedhoff HR, Chen S, Balderrama A, Sun GY, Koopmans B, DePalma RG, et al. Long-Term Effects of Low-Intensity Blast Non-Inertial Brain Injury on Anxiety-Like Behaviors in Mice: Home-Cage Monitoring Assessments. Neurotrauma Rep. 2022;3(1):27–38. Epub 20220111. doi: 10.1089/neur.2021.0063. PubMed PMID: 35141713; PubMed Central PMCID: PMCPMC8820222.

55. Angelakos CC, Tudor JC, Ferri SL, Jongens TA, Abel T. Home-cage hypoactivity in mouse genetic models of autism spectrum disorder. Neurobiol Learn Mem. 2019;165:107000. Epub 20190220. doi: 10.1016/j.nlm.2019.02.010. PubMed PMID: 30797034; PubMed Central PMCID: PMCPMC6913530.

56. Qu W, Liu NK, Xie XM, Li R, Xu XM. Automated monitoring of early neurobehavioral changes in mice following traumatic brain injury. Neural Regen Res. 2016;11(2):248–56. doi: 10.4103/1673-5374.177732. PubMed PMID: 27073377; PubMed Central PMCID: PMCPMC4810988.

57. Robinson-Junker AL, O’Hara B F, Gaskill BN. Out Like a Light? The Effects of a Diurnal Husbandry Schedule on Mouse Sleep and Behavior. J Am Assoc Lab Anim Sci. 2018;57(2):124–33. Epub 2018/03/21. PubMed PMID: 29555001; PubMed Central PMCID: PMCPMC5868378.

58. Tang X, Liu X, Yang L, Sanford LD. Rat strain differences in sleep after acute mild stressors and short-term sleep loss. Behav Brain Res. 2005;160(1):60–71. Epub 2005/04/20. doi: 10.1016/j.bbr.2004.11.015. PubMed PMID: 15836901.

59. Urban R, Scherrer G, Goulding EH, Tecott LH, Basbaum AI. Behavioral indices of ongoing pain are largely unchanged in male mice with tissue or nerve injury-induced mechanical hypersensitivity. Pain. 2011;152(5):990–1000. Epub 20110121. doi: 10.1016/j.pain.2010.12.003. PubMed PMID: 21256675; PubMed Central PMCID: PMCPMC3079194.

60. Shenk J, Lohkamp KJ, Wiesmann M, Kiliaan AJ. Automated Analysis of Stroke Mouse Trajectory Data With Traja. Front Neurosci. 2020;14:518. Epub 2020/06/12. doi: 10.3389/fnins.2020.00518. PubMed PMID: 32523509; PubMed Central PMCID: PMCPMC7262161.

61. Pritchett D, Jagannath A, Brown LA, Tam SKE, Hasan S, Gatti S, et al. Deletion of Metabotropic Glutamate Receptors 2 and 3 (mGlu2 & mGlu3) in Mice Disrupts Sleep and Wheel-Running Activity, and Increases the Sensitivity of the Circadian System to Light. PLOS ONE. 2015;10(5):e0125523. doi: 10.1371/journal.pone.0125523.

62. Júdice PB, Hetherington-Rauth M, Northstone K, Andersen LB, Wedderkopp N, Ekelund U, et al. Changes in Physical Activity and Sedentary Patterns on Cardiometabolic Outcomes in the Transition to Adolescence: International Children’s Accelerometry Database 2.0. J Pediatr. 2020. Epub 2020/06/20. doi: 10.1016/j.jpeds.2020.06.018. PubMed PMID: 32553870.

63. Varma VR, Dey D, Leroux A, Di J, Urbanek J, Xiao L, et al. Re-evaluating the effect of age on physical activity over the lifespan. Prev Med. 2017;101:102–8. Epub 2017/06/06. doi: 10.1016/j.ypmed.2017.05.030. PubMed PMID: 28579498; PubMed Central PMCID: PMCPMC5541765.

64. Oikawa SY, Holloway TM, Phillips SM. The Impact of Step Reduction on Muscle Health in Aging: Protein and Exercise as Countermeasures. Front Nutr. 2019;6:75. Epub 2019/06/11. doi: 10.3389/fnut.2019.00075. PubMed PMID: 31179284; PubMed Central PMCID: PMCPMC6543894.

65. Palmberg L, Rantalainen T, Rantakokko M, Karavirta L, Siltanen S, Skantz H, et al. The Associations of Activity Fragmentation with Physical and Mental Fatigability among Community-Dwelling 75-, 80- and 85-Year-Old People. J Gerontol A Biol Sci Med Sci. 2020. Epub 2020/07/03. doi: 10.1093/gerona/glaa166. PubMed PMID: 32614396.

66. Smirnova E, Leroux A, Cao Q, Tabacu L, Zipunnikov V, Crainiceanu C, et al. The Predictive Performance of Objective Measures of Physical Activity Derived From Accelerometry Data for 5-Year All-Cause Mortality in Older Adults: National Health and Nutritional Examination Survey 2003-2006. J Gerontol A Biol Sci Med Sci. 2020;75(9):1779–85. Epub 2019/09/11. doi: 10.1093/gerona/glz193. PubMed PMID: 31504213; PubMed Central PMCID: PMCPMC7494021.

67. Štefelová N, Dygrýn J, Hron K, Gába A, Rubín L, Palarea-Albaladejo J. Robust Compositional Analysis of Physical Activity and Sedentary Behaviour Data. Int J Environ Res Public Health. 2018;15(10). Epub 2018/10/17. doi: 10.3390/ijerph15102248. PubMed PMID: 30322203; PubMed Central PMCID: PMCPMC6210094.

68. Hamer M, Stamatakis E, Chastin S, Pearson N, Brown M, Gilbert E, et al. Feasibility of Measuring Sedentary Time Using Data From a Thigh-Worn Accelerometer. Am J Epidemiol. 2020;189(9):963–71. Epub 2020/03/29. doi: 10.1093/aje/kwaa047. PubMed PMID: 32219368; PubMed Central PMCID: PMCPMC7443760.

69. Fishman EI, Steeves JA, Zipunnikov V, Koster A, Berrigan D, Harris TA, et al. Association between Objectively Measured Physical Activity and Mortality in NHANES. Med Sci Sports Exerc. 2016;48(7):1303–11. Epub 2016/02/06. doi: 10.1249/mss.0000000000000885. PubMed PMID: 26848889; PubMed Central PMCID: PMCPMC4911242.

70. Nastasi AJ, Ahuja A, Zipunnikov V, Simonsick EM, Ferrucci L, Schrack JA. Objectively Measured Physical Activity and Falls in Well-Functioning Older Adults: Findings From the Baltimore Longitudinal Study of Aging. Am J Phys Med Rehabil. 2018;97(4):255–60. Epub 2017/09/16. doi: 10.1097/phm.0000000000000830. PubMed PMID: 28915202; PubMed Central PMCID: PMCPMC5851797.

71. Wanigatunga AA, D. J, Zipunnikov V, Urbanek JK, Kuo PL, Simonsick EM, et al. Association of Total Daily Physical Activity and Fragmented Physical Activity With Mortality in Older Adults. JAMA Netw Open. 2019;2(10):e1912352. Epub 2019/10/03. doi: 10.1001/jamanetworkopen.2019.12352. PubMed PMID: 31577355; PubMed Central PMCID: PMCPMC6777397.

72. Merikangas KR, Swendsen J, Hickie IB, Cui L, Shou H, Merikangas AK, et al. Real-time Mobile Monitoring of the Dynamic Associations Among Motor Activity, Energy, Mood, and Sleep in Adults With Bipolar Disorder. JAMA Psychiatry. 2019;76(2):190–8. Epub 2018/12/13. doi: 10.1001/jamapsychiatry.2018.3546. PubMed PMID: 30540352; PubMed Central PMCID: PMCPMC6439734 commissioner of Australia’s National Mental Health Commission; being a codirector of Health and Policy at the Brain and Mind Centre, University of Sydney; leading community-based and pharmaceutical industry–supported (Wyeth, Eli Lily, Servier, Pfizer, and AstraZeneca) projects focused on the identification and better management of anxiety and depression; being a member of the Medical Advisory Panel for Medibank Private until October 2017; being a board member of Psychosis Australia Trust and a member of the Veterans Mental Health Clinical Reference group; and being the Chief Scientific Advisor to and an equity shareholder in Innowell. No other disclosures were reported.

73. Schrack JA, Kuo PL, Wanigatunga AA, D. J, Simonsick EM, Spira AP, et al. Active-to-Sedentary Behavior Transitions, Fatigability, and Physical Functioning in Older Adults. J Gerontol A Biol Sci Med Sci. 2019;74(4):560–7. Epub 2018/10/26. doi: 10.1093/gerona/gly243. PubMed PMID: 30357322; PubMed Central PMCID: PMCPMC6417447.

74. Lamers F, Swendsen J, Cui L, Husky M, Johns J, Zipunnikov V, et al. Mood reactivity and affective dynamics in mood and anxiety disorders. J Abnorm Psychol. 2018;127(7):659–69. Epub 2018/10/20. doi: 10.1037/abn0000378. PubMed PMID: 30335438.

75. Melin M, Hagerman I, Gonon A, Gustafsson T, Rullman E. Variability in Physical Activity Assessed with Accelerometer Is an Independent Predictor of Mortality in CHF Patients. PLoS One. 2016;11(4):e0153036. Epub 2016/04/08. doi: 10.1371/journal.pone.0153036. PubMed PMID: 27054323; PubMed Central PMCID: PMCPMC4824362.

76. Forkosh O, Karamihalev S, Roeh S, Alon U, Anpilov S, Touma C, et al. Identity domains capture individual differences from across the behavioral repertoire. Nat Neurosci. 2019;22(12):2023–8. Epub 2019/11/07. doi: 10.1038/s41593-019-0516-y. PubMed PMID: 31686022.

77. Cain KL, Sallis JF, Conway TL, Van Dyck D, Calhoon L. Using accelerometers in youth physical activity studies: a review of methods. J Phys Act Health. 2013;10(3):437–50. Epub 2013/04/27. doi: 10.1123/jpah.10.3.437. PubMed PMID: 23620392; PubMed Central PMCID: PMCPMC6331211.

78. Bao W, Sun Y, Zhang T, Zou L, Wu X, Wang D, et al. Exercise Programs for Muscle Mass, Muscle Strength and Physical Performance in Older Adults with Sarcopenia: A Systematic Review and Meta-Analysis. Aging Dis. 2020;11(4):863–73. Epub 2020/08/09. doi: 10.14336/ad.2019.1012. PubMed PMID: 32765951; PubMed Central PMCID: PMCPMC7390512.

79. Varma VR, Dey D, Leroux A, Di J, Urbanek J, Xiao L, et al. Total volume of physical activity: TAC, TLAC or TAC(λ). Prev Med. 2018;106:233–5. Epub 2017/10/31. doi: 10.1016/j.ypmed.2017.10.028. PubMed PMID: 29080825; PubMed Central PMCID: PMCPMC5897896.

80. Byrom B, Stratton G, Mc Carthy M, Muehlhausen W. Objective measurement of sedentary behaviour using accelerometers. Int J Obes (Lond). 2016;40(11):1809–12. Epub 2016/08/02. doi: 10.1038/ijo.2016.136. PubMed PMID: 27478922; PubMed Central PMCID: PMCPMC5116050.

81. Godfrey A, Vandendriessche B, Bakker JP, Fitzer-Attas C, Gujar N, Hobbs M, et al. Fit-for-Purpose Biometric Monitoring Technologies: Leveraging the Laboratory Biomarker Experience. Clin Transl Sci. 2021;14(1):62–74. Epub 2020/08/10. doi: 10.1111/cts.12865. PubMed PMID: 32770726; PubMed Central PMCID: PMCPMC7877826 owns company stock; J.P.B is a full-time employee at Philips; N.G. is a full-time employee at Samsung NeuroLogica. E.S.I. is an employee of Koneksa Health and owns company stock; C.A.N. is an employee of Pfizer, Inc. and owns company stock; V.P. is a full-time employee at Takeda; W.A.W. is an Advisor of Koneksa and Elektra Labs, a consultant for Best Doctors/Teladoc and has research funding from Genentech and Pfizer; C.F.A. is a full-time employee at Mitsubishi Tanabe Pharma America. All other authors declared no competing interests for this work. As Editor-in-Chief of Clinical & Translational Science, John A. Wagner was not involved in the review or decision process for this paper.

82. Karas M, Bai J, Strączkiewicz M, Harezlak J, Glynn NW, Harris T, et al. Accelerometry data in health research: challenges and opportunities. Stat Biosci. 2019;11(2):210–37. Epub 2019/11/26. doi: 10.1007/s12561-018-9227-2. PubMed PMID: 31762829; PubMed Central PMCID: PMCPMC6874221.

83. Baran SW, Lim MA, Do JP, Stolyar P, Rabe MD, Schaevitz LR, et al. Digital biomarkers enable automated, longitudinal monitoring in a mouse model of aging. J Gerontol A Biol Sci Med Sci. 2021. Epub 2021/01/26. doi: 10.1093/gerona/glab024. PubMed PMID: 33491048.

84. Volker LA, Maar BA, Pulido Guevara BA, Bilkei-Gorzo A, Zimmer A, Bronneke H, et al. Neph2/Kirrel3 regulates sensory input, motor coordination, and home-cage activity in rodents. Genes Brain Behav. 2018;17(8):e12516. Epub 2018/08/23. doi: 10.1111/gbb.12516. PubMed PMID: 30133126.

85. Loh DH, Kudo T, Truong D, Wu Y, Colwell CS. The Q175 Mouse Model of Huntington’s Disease Shows Gene Dosage- and Age-Related Decline in Circadian Rhythms of Activity and Sleep. PLOS ONE. 2013;8(7):e69993. doi: 10.1371/journal.pone.0069993.

86. Bonasera SJ, Chaudoin TR, Goulding EH, Mittek M, Dunaevsky A. Decreased home cage movement and oromotor impairments in adult Fmr1-KO mice. Genes Brain Behav. 2017;16(5):564–73. Epub 2017/02/22. doi: 10.1111/gbb.12374. PubMed PMID: 28218824; PubMed Central PMCID: PMCPMC6042514.

87. Hasriadi, Dasuni Wasana PW, Vajragupta O, Rojsitthisak P, Towiwat P. Automated home-cage monitoring as a potential measure of sickness behaviors and pain-like behaviors in LPS-treated mice. PLoS One. 2021;16(8):e0256706. Epub 20210827. doi: 10.1371/journal.pone.0256706. PubMed PMID: 34449819; PubMed Central PMCID: PMCPMC8396795.

88. Lim MA, Defensor EB, Mechanic JA, Shah PP, Jaime EA, Roberts CR, et al. Retrospective Analysis of the Effects of Identification Procedures and Cage Changing by Using Data from Automated, Continuous Monitoring. J Am Assoc Lab Anim Sci. 2019;58(2):126–41. Epub 20190214. doi: 10.30802/aalas-jaalas-18-000056. PubMed PMID: 30764898; PubMed Central PMCID: PMCPMC6433355.

89. Marcus AD, Achanta S, Jordt SE. Protocol for non-invasive assessment of spontaneous movements of group-housed animals using remote video monitoring. STAR Protoc. 2022;3(2):101326. Epub 20220414. doi: 10.1016/j.xpro.2022.101326. PubMed PMID: 35479115; PubMed Central PMCID: PMCPMC9036393.

90. Balcombe JP. Laboratory environments and rodents’ behavioural needs: a review. Lab Anim. 2006;40(3):217–35. Epub 2006/06/29. doi: 10.1258/002367706777611488. PubMed PMID: 16803640.

91. Liu N, Wang Y, An AY, Banker C, Qian YH, O’Donnell JM. Single housing-induced effects on cognitive impairment and depression-like behavior in male and female mice involve neuroplasticity-related signaling. Eur J Neurosci. 2020;52(1):2694–704. Epub 2019/09/01. doi: 10.1111/ejn.14565. PubMed PMID: 31471985.

92. Škop V, Xiao C, Liu N, Gavrilova O, Reitman ML. The effects of housing density on mouse thermal physiology depend on sex and ambient temperature. Mol Metab. 2021;53:101332. Epub 2021/09/04. doi: 10.1016/j.molmet.2021.101332. PubMed PMID: 34478905; PubMed Central PMCID: PMCPMC8463779.

93. Hebda-Bauer EK, Dokas LA, Watson SJ, Akil H. Adaptation to single housing is dynamic: Changes in hormone levels, gene expression, signaling in the brain, and anxiety-like behavior in adult male C57Bl/6J mice. Horm Behav. 2019;114:104541. Epub 2019/06/21. doi: 10.1016/j.yhbeh.2019.06.005. PubMed PMID: 31220462; PubMed Central PMCID: PMCPMC7466935.

94. Nagy TR, Krzywanski D, Li J, Meleth S, Desmond R. Effect of group vs. single housing on phenotypic variance in C57BL/6J mice. Obes Res. 2002;10(5):412–5. Epub 2002/05/15. doi: 10.1038/oby.2002.57. PubMed PMID: 12006642.

95. Kappel S, Hawkins P, Mendl MT. To Group or Not to Group? Good Practice for Housing Male Laboratory Mice. Animals (Basel). 2017;7(12). Epub 20171124. doi: 10.3390/ani7120088. PubMed PMID: 29186765; PubMed Central PMCID: PMCPMC5742782.

96. Charlier R, Knaeps S, Mertens E, Van Roie E, Delecluse C, Lefevre J, et al. Age-related decline in muscle mass and muscle function in Flemish Caucasians: a 10-year follow-up. Age (Dordr). 2016;38(2):36. Epub 2016/03/11. doi: 10.1007/s11357-016-9900-7. PubMed PMID: 26961694; PubMed Central PMCID: PMCPMC5005902.

97. Clemenza K, Weiss SH, Cheslack K, Kandel DB, Kandel ER, Levine AA. Social isolation is closely linked to a marked reduction in physical activity in male mice. J Neurosci Res. 2021;99(4):1099–107. Epub 20201226. doi: 10.1002/jnr.24777. PubMed PMID: 33368537.

98. Jirkof P, Cesarovic N, Rettich A, Fleischmann T, Arras M. Individual housing of female mice: influence on postsurgical behaviour and recovery. Lab Anim. 2012;46(4):325–34. doi: 10.1258/la.2012.012027. PubMed PMID: 23097566.

99. Mertens S, Vogt M, Gass P, Palme R, Hiebl B, Chourbaji S. Effect of three different forms of handling on the variation of aggression-associated parameters in individually and group-housed male C57BL/6NCrl mice. PLOS ONE. 2019;14:e0215367. doi: 10.1371/journal.pone.0215367.

100. Kamakura R, Kovalainen M, Leppäluoto J, Herzig K-H, Mäkelä KA. The effects of group and single housing and automated animal monitoring on urinary corticosterone levels in male C57BL/6 mice. Physiological reports. 2016;4(3):e12703. doi: 10.14814/phy2.12703. PubMed PMID: 26869685.

101. Hunt C, Hambly C. Faecal corticosterone concentrations indicate that separately housed male mice are not more stressed than group housed males. Physiol Behav. 2006;87(3):519–26. Epub 20060125. doi: 10.1016/j.physbeh.2005.11.013. PubMed PMID: 16442135.

102. Arndt SS, Laarakker MC, van Lith HA, van der Staay FJ, Gieling E, Salomons AR, et al. Individual housing of mice--impact on behaviour and stress responses. Physiol Behav. 2009;97(3-4):385–93. Epub 20090318. doi: 10.1016/j.physbeh.2009.03.008. PubMed PMID: 19303031.

