## Supporting information for "Bouts of rest and physical activity in C57BL/6J mice"

### Supportive Information

#### Table S1

Data sequencies removed from the original files and not included in the results or statistical tests. For further information see Material and methods section*.*

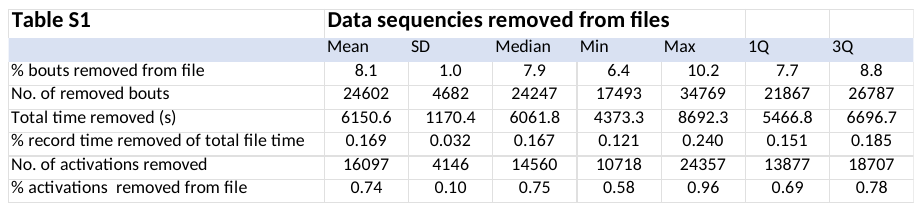

#### Fig. S1

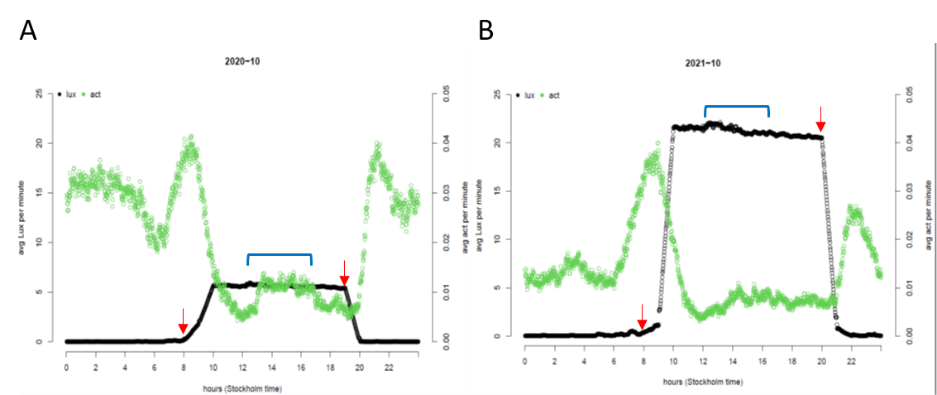

**Fig. S1** Panel shows light on (left red arrow in each panel) and lights off (red arrow to the right). Abscissa is local time and left ordinate Lux at the rack front. Right ordinate shows EAD min^-1^ across the LD cycle for Fx3 cohort used in this study. Blue bracket indicates time for cage change and the immediate increase in activity that follows. Note that the response in activity precedes lights on, peaks during the dawn period and decreases towards the end of dawn. Dusk is followed by an increase in EAD, the ramp up occurs during the first hour following dusk initiation. Data on Lux were recorded by the REM unit on the DVC rack, EAD by the DVC.

#### Fig S2

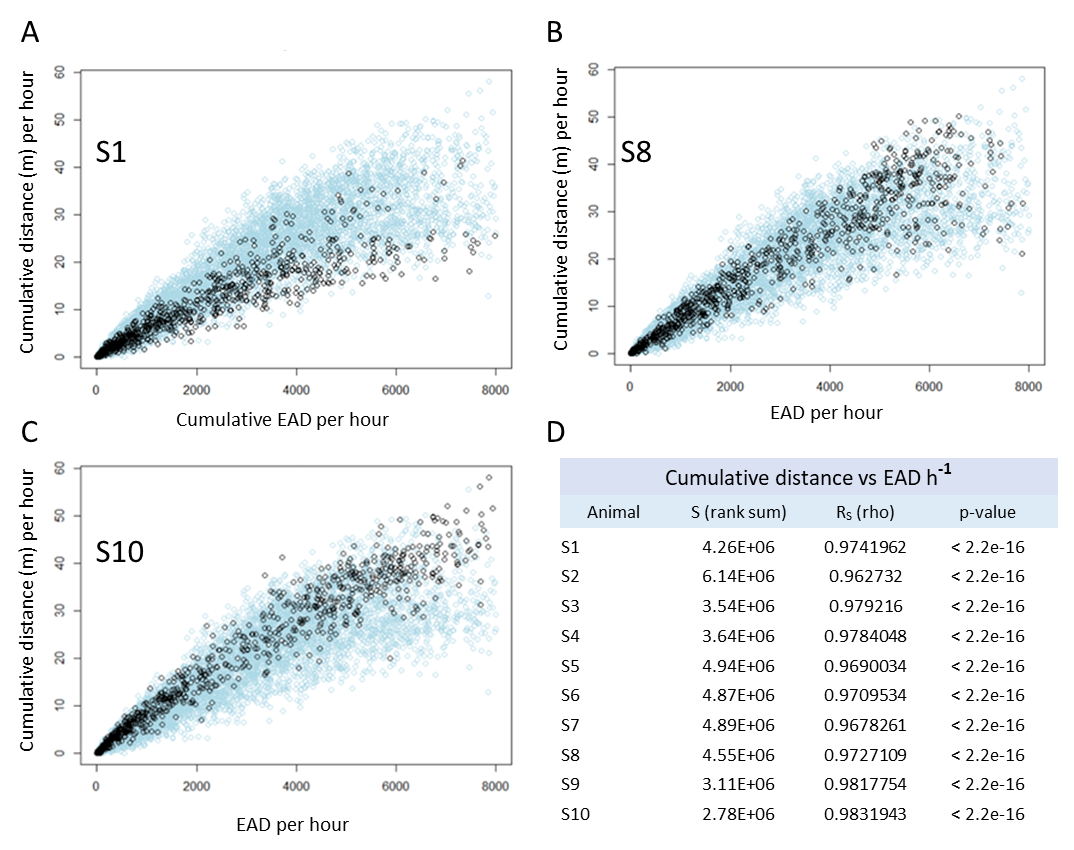

**Fig S2** A-C Plots showing the correlation between distance travelled per hour (the tracking metric, ordinate) and number of electrode activations (EAD metric, abscissa) for three of the single-housed (S1, S8 and S10) mice (black circles). Each data set was plotted on the whole data set of 10 mice (green circles). In D, the Spearman rank correlation data for all 10 mice have been compiled.

#### Fig S3

EAD min-1

|  | Females | Males |
| --- | --- | --- |
| -5 | 0.0273229 | 0.0150185 |
| -4 | 0.0167169 | 0.0116641 |
| -3 | 0.0166544 | 0.00846056 |
| -2 | 0.00542512 | 0.0042451 |
| -1 | 0.00511232 | 0.00346657 |
| 0 | 9.91776e-06 | 0.000102421 |
| 0 | 8.01421e-07 | 6.31756e-06 |
| 1 | 0.0101829 | 0.00609844 |
| 2 | 0.0122561 | 0.00976832 |
| 3 | 0.010305 | 0.00934523 |
| 4 | 0.0179349 | 0.0124729 |
| 5 | 0.0316016 | 0.0196842 |

**Fig S3** Histogram showing the impact on EAD min^-1^ by stepwise lowering the housing density from 5 to 1 mice (one mouse was removed each consecutive day) and then adding the mice back one by one up to five. Female cages (n=3) in red and male cages (n=5) in blue; average EAD value min-1 across 24 h (λ=2) with standard deviation has been indicated.

#### Fig S4

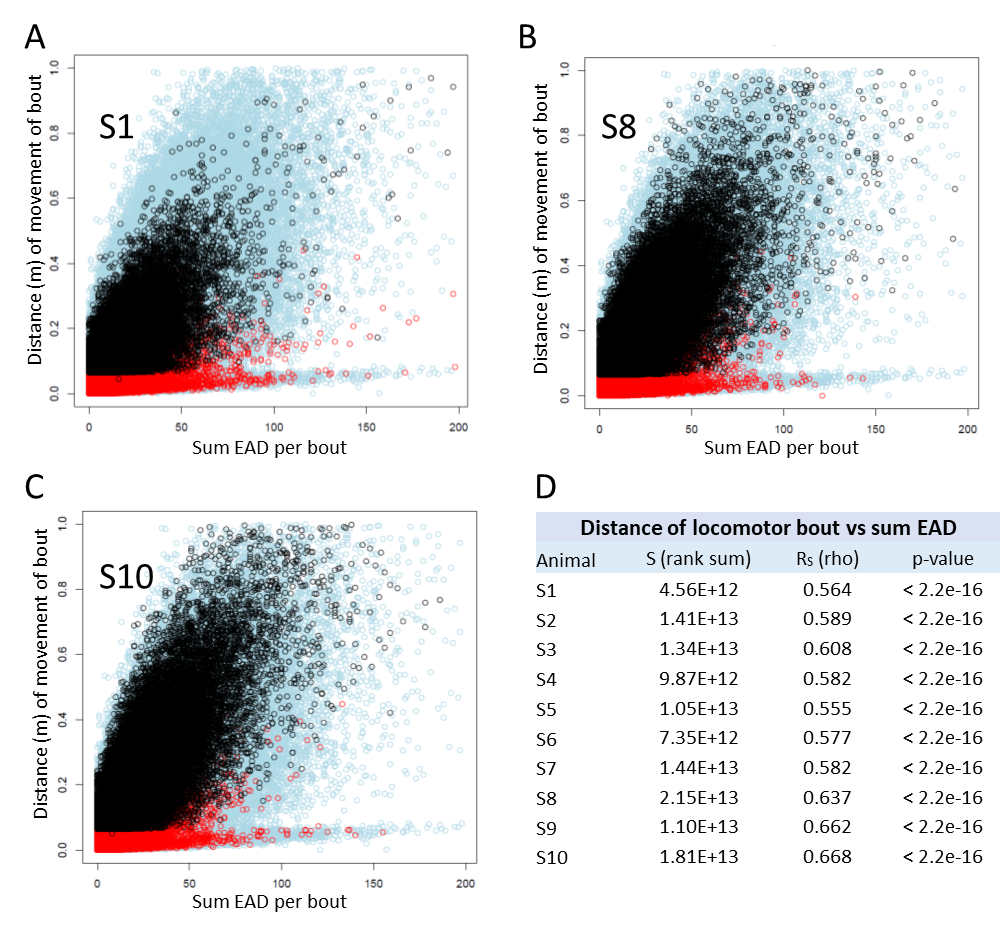

**Fig S4** A-C Plots showing the correlation between distance travelled per bout (the tracking metric, ordinate) and number of electrode activations per bout (EAD metric, abscissa) for three (S1, S8 and S10) of the single-housed mice. Locomotor bouts have been indicated by black circles and MOTS bouts by red circles. Data for each of the three mice was plotted on the whole data set from 10 mice (green circles). In D, the Spearman rank correlation data for locomotor bouts vs EAD for all 10 mice have been compiled.

#### Fig S5

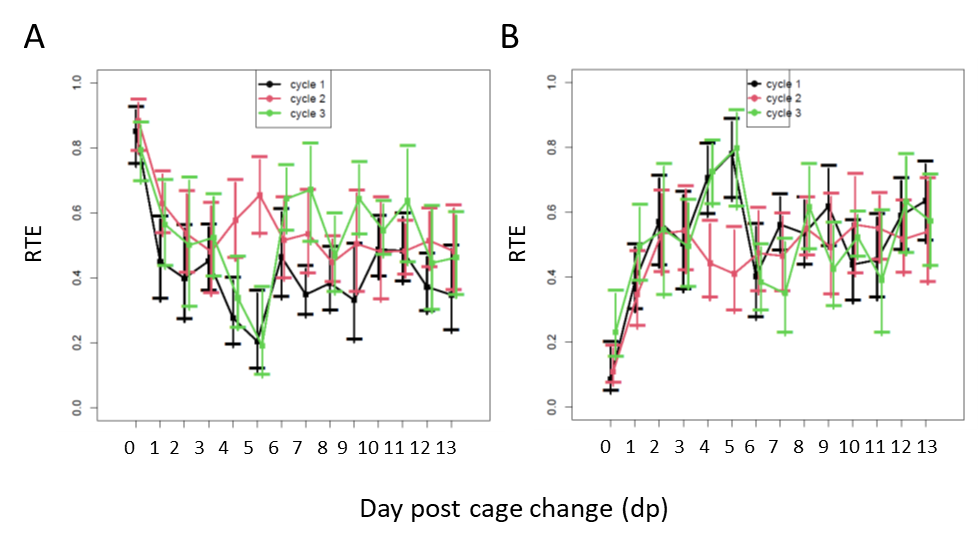

**Fig S5** A-B show the relative effect size of days-post-cage-change (dp) and cage-change-cycle (CC, colour coded in black, red and green), respectively. In A (model: PA ~dp * CC, the major impact is by dp (p=2E-9) and only little by CC (p=0.02). D shows that CC has no significant impact on long rest (p=0.51) while dp has a strong effect (p=3.8E-11).

#### Fig S6

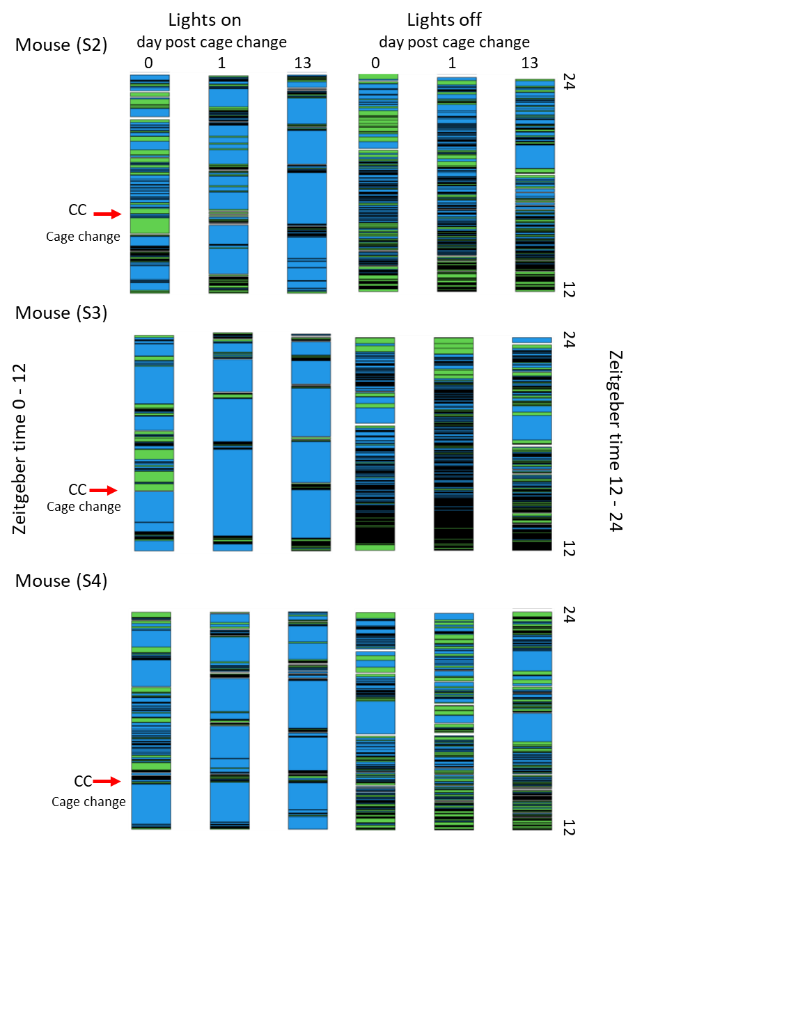

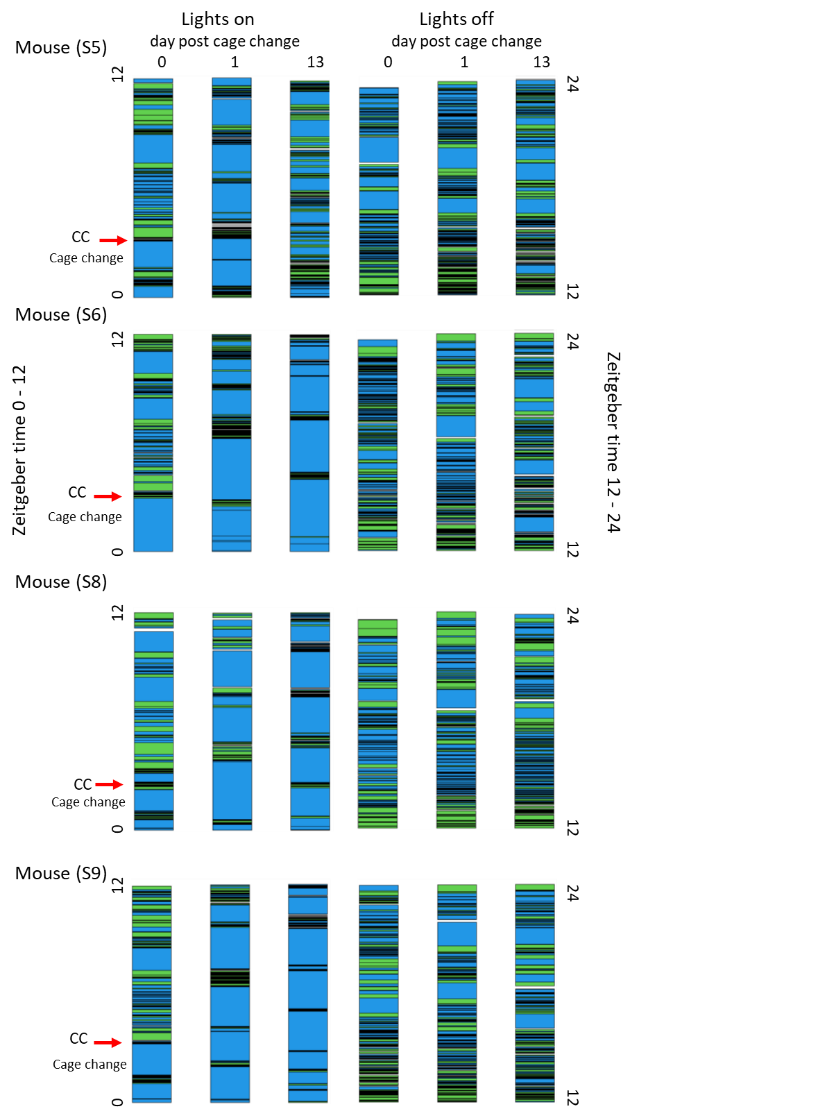

**Fig S6** Pattern of long-rest episodes during lights on (ZT 0-12) and lights off (ZT 12-24) (DL 12:12) for the mice S2-S6 and S8-S9 mice housed in isolation. Long rest bouts indicated by blue colour. Bouts disrupting long-rest periods in black/green. Left ordinate is 12h lights on to the left and 12h lights off on the ordinate to the right. Columns are day post cage change (dp) where 0 is the day of the cage change (CC; red arrow indicates the time for CC) for lights on (to the left) and lights off (to the right). Dp1 the day after CC and dp13 the day before the next CC.

#### Fig. S7

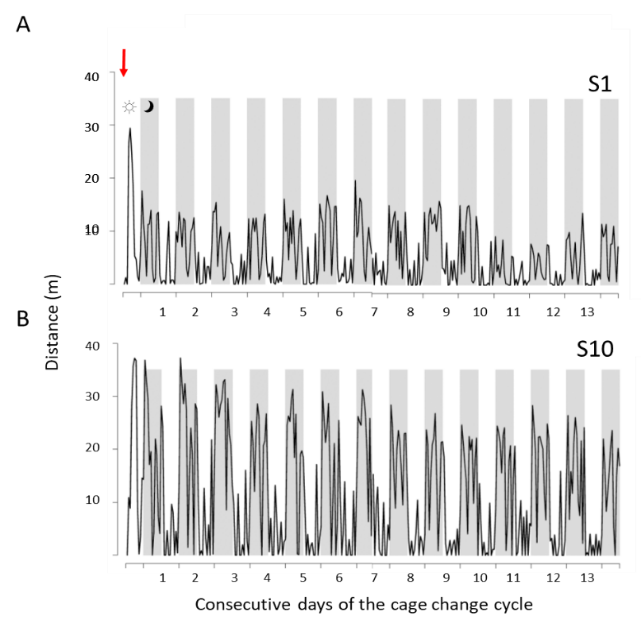

**Fig S7** A-B show mouse S1 and S10 distance per hour (m; ordinate) made through the LD cycle of each consecutive day (abscissa) of a cage change cycle. Time of the cage change marked with red arrow in top panel. L (ZT 0-12) on white background, while D (ZT 12-24) have a grey background; see symbols in upper panel.

#### Fig. S8

S3	S4	S5	S6

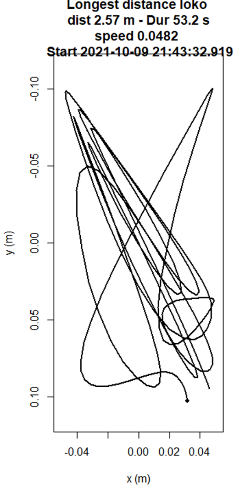

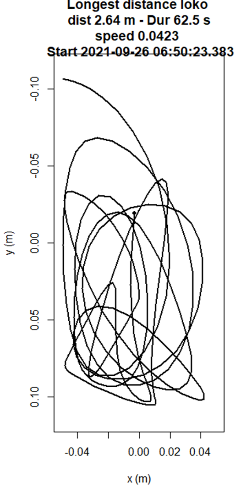

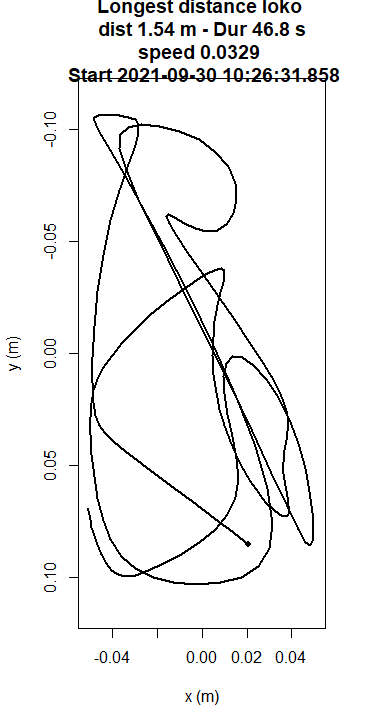

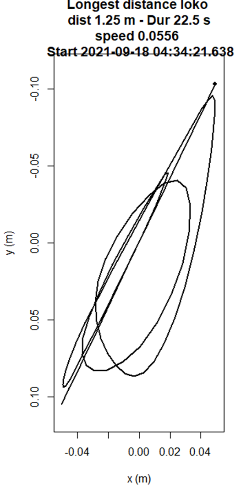

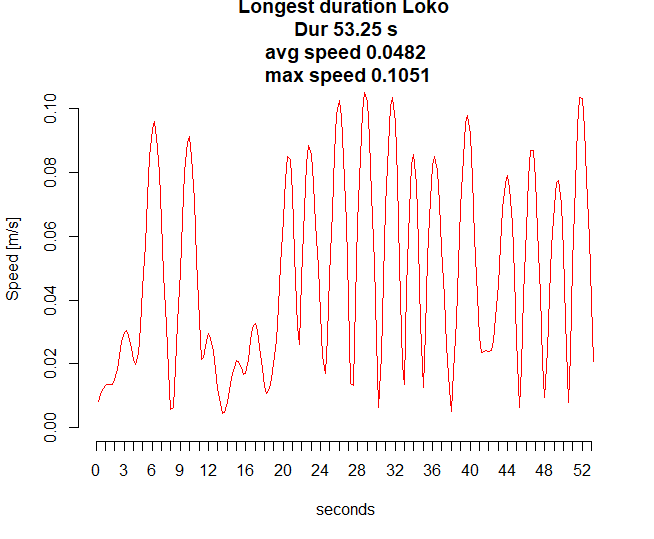

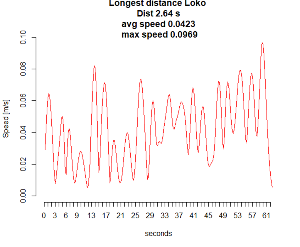

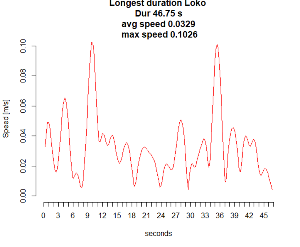

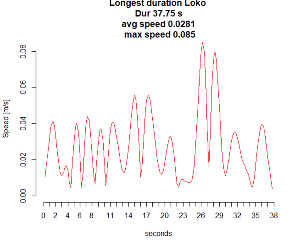

S7	S8	S9

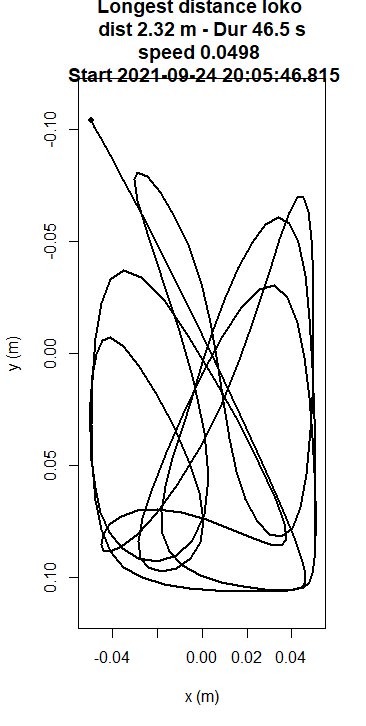

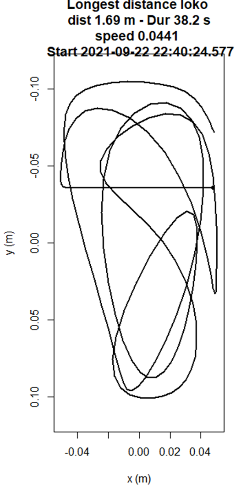

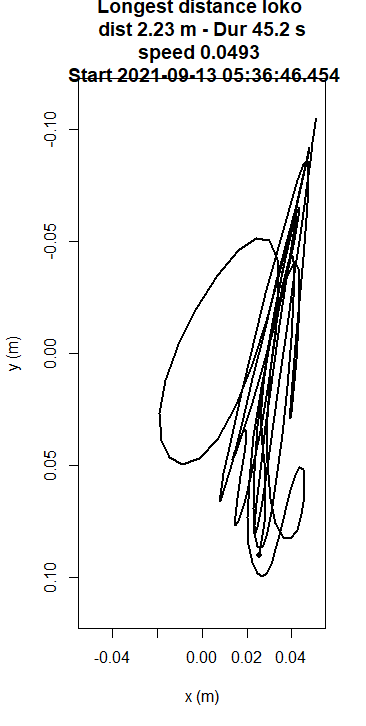

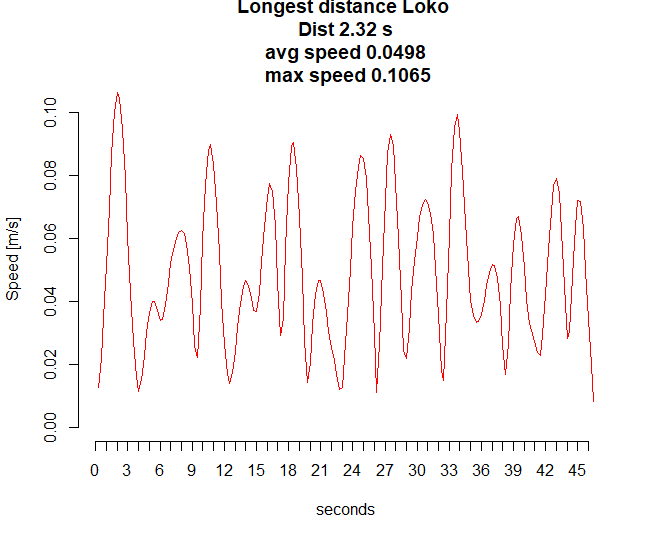

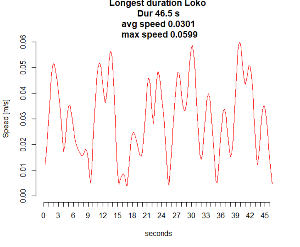

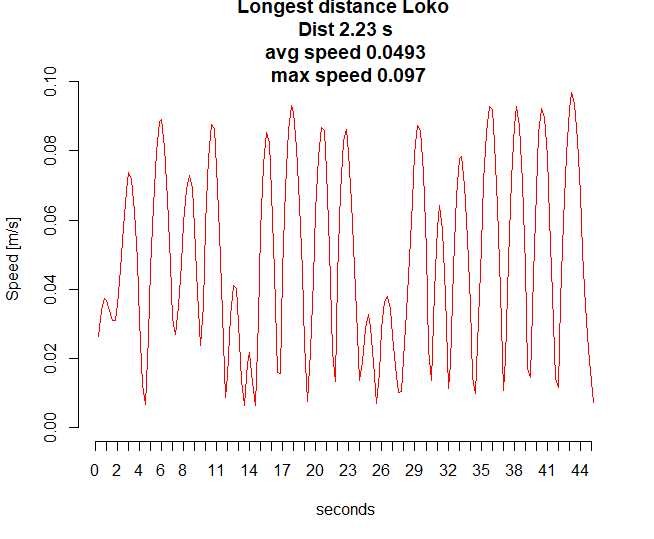

**Fig S8** Panels show longest recorded locomotor bout for mouse S3-S9, respectively. Lower panels with red traces show corresponding speedograms. Plots and speedograms by the trajaR software.

#### Fig S9

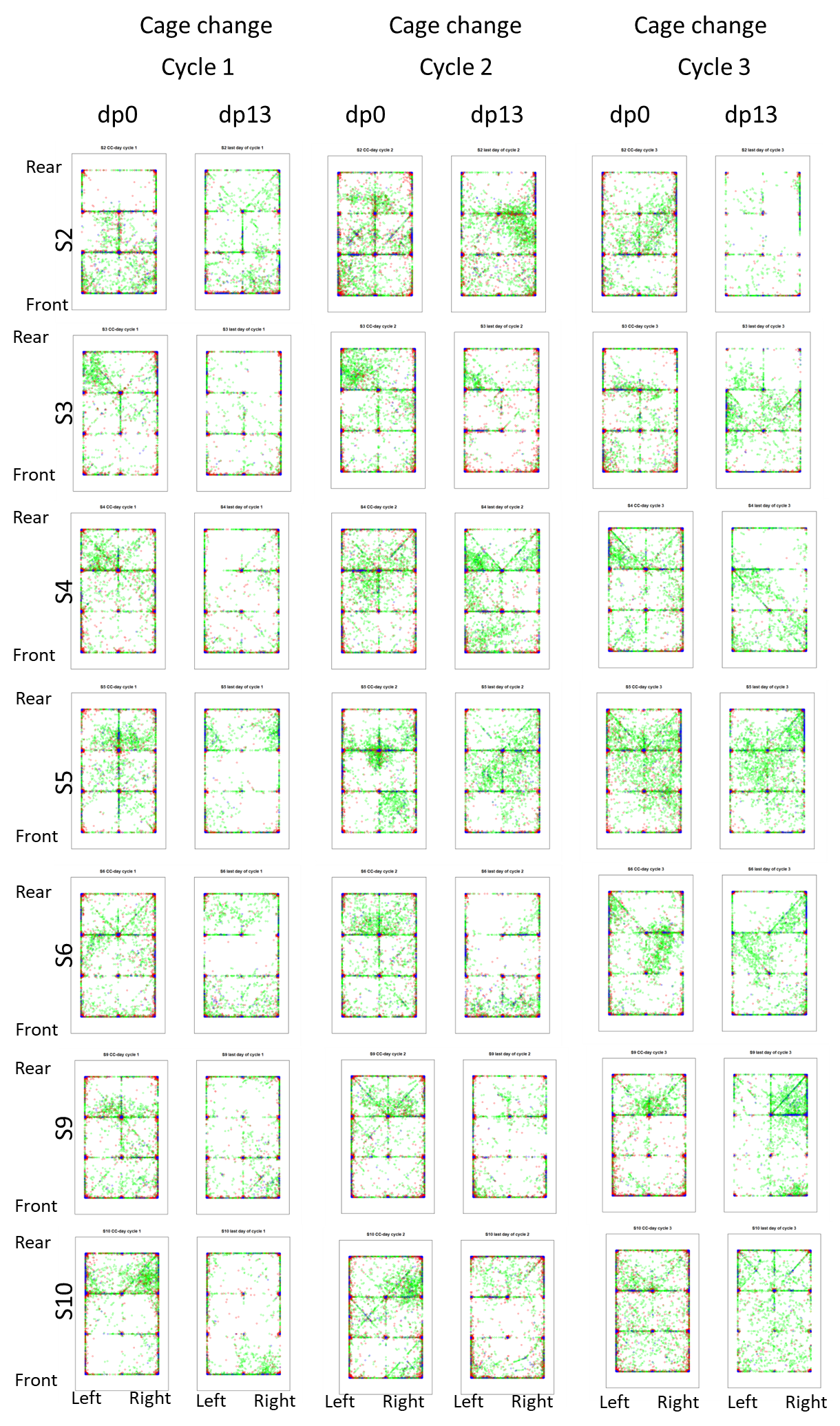

**Fig S9** Panels show from left to right the cage floor with rear up and front down at the day of the cage-change (dp0) and the final day prior to the next cage-change (dp13) in three consecutive cage change cycles. In each cage floor panel, the starting point (x, y coordinates) for each bout of that day as been indicated. Bouts of locomotion is indicated in red, long rest in blue, short rest in orange and MOTS in green. Each row of panels is from one animal (S2, S3 etc.).

#### Fig. S10

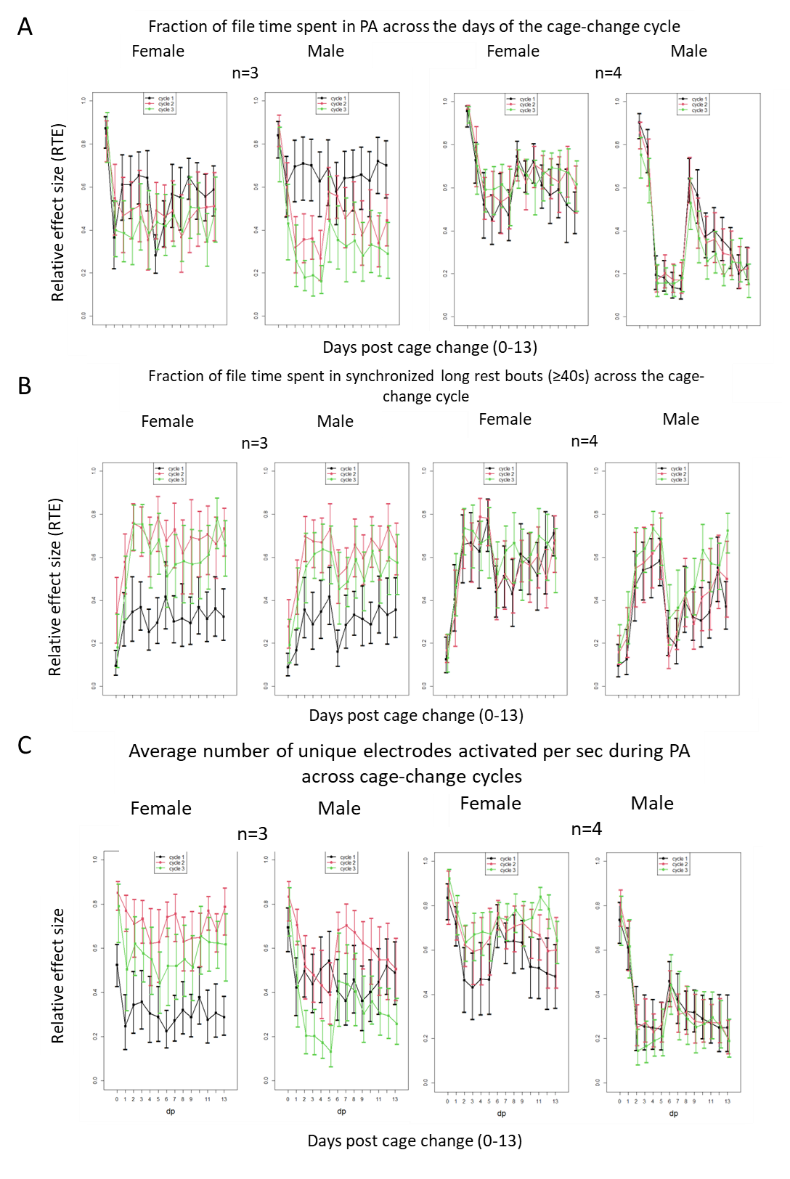

**Fig. S10** A shows the relative effect size (RTE; ordinate) on PA of sex, dp (0-13; abscissa) and cage change cycle (CC; colour coded black, red, and green in each panel) have been indicated at densities x3 and x4 for female and male mice.

B shows the relative effect size (RTE; ordinate) on long rest of sex, dp (0-13; abscissa) and cage change cycle (CC; colour coded black, red, and green in each panel) have been indicated at densities x3 and x4. Dp had the strongest impact on time in long rest, followed by sex at density n=4

C shows the relative effect size (RTE; ordinate) on number of unique electrode activations s^-1^ of sex, dp (0-13; abscissa) and cage-change cycle (CC; colour coded in each panel; black, red, and green) have been indicated.
